# Proteome asymmetry in mouse and human embryos before fate specification

**DOI:** 10.1101/2024.08.26.609777

**Authors:** Lisa K. Iwamoto-Stohl, Aleksandra A. Petelski, Maciej Meglicki, Audrey Fu, Saad Khan, Harrison Specht, Gray Huffman, Jason Derks, Victoria Jorgensen, Bailey A.T. Weatherbee, Antonia Weberling, Carlos W. Gantner, Rachel S. Mandelbaum, Richard J. Paulson, Lisa Lam, Ali Ahmady, Estefania Sanchez Vasquez, Nikolai Slavov, Magdalena Zernicka-Goetz

**Affiliations:** University of Cambridge, Department of Physiology, Development and Neuroscience; Downing Street, Cambridge, CB2 3DY, UK; California Institute of Technology (Caltech), Division of Biology and Biological Engineering, Pasadena, CA 91125, USA; Northeastern University, Department of Bioengineering and Barnett Institute, Boston, MA 02115, USA; Parallel Squared Technology Institute, Watertown, MA, USA; University of Cambridge, Department of Biochemistry, Hopkins Building, Tennis Court Road, Cambridge CB2 1QW, UK; Division of Reproductive Endocrinology and Infertility, Department of Obstetrics and Gynecology, University of Southern California, Los Angeles, California, USA; Division of Reproductive Endocrinology and Infertility, Department of Obstetrics and Gynecology, Washington University in St. Louis, St. Louis, Missouri, USA

## Abstract

Pre-patterning of the embryo, driven by spatially localized factors, is a common feature across several non-mammalian species^1–4^. However, mammals display regulative development and thus it was thought that blastomeres of the embryo do not show such pre-patterning, contributing randomly to the three lineages of the blastocyst: the epiblast, primitive endoderm and trophectoderm that will generate the new organism, the yolk sac and placenta respectively ^4–6^. Unexpectedly, early blastomeres of mouse and human embryos have been reported to have distinct developmental fates, potential and heterogeneous abundance of certain transcripts^7–12^. Nevertheless, the extent of the earliest intra-embryo differences remains unclear and controversial. Here, by utilizing multiplexed and label-free single-cell proteomics by mass-spectrometry^13^, we show that 2-cell mouse and human embryos contain an alpha and a beta blastomere as defined by differential abundance of hundreds of proteins exhibiting strong functional enrichment for protein synthesis, transport, and degradation. Such asymmetrically distributed proteins include Gps1 and Nedd8, depletion or overexpression of which in one blastomere of the 2-cell embryo impacts lineage segregation. These protein asymmetries increase at 4-cell stage. Intriguingly, halved mouse zygotes display asymmetric protein abundance that resembles alpha and beta blastomeres, suggesting differential proteome localization already within zygotes. We find that beta blastomeres give rise to a blastocyst with a higher proportion of epiblast cells than alpha blastomeres and that vegetal blastomeres, which are known to have a reduced developmental potential, are more likely to be alpha. Human 2-cell blastomeres also partition into two clusters sharing strong concordance with clusters found in mouse, in terms of differentially abundant proteins and functional enrichment. To our knowledge, this is the first demonstration of intra-zygotic and inter-blastomere proteomic asymmetry in mammals that has a role in lineage segregation.

## Main

In mammals, including mouse and human, the fertilized egg undergoes cleavage divisions to give rise to embryonic and extraembryonic cell types^14,15^. Two cell fate decisions are crucial for blastocyst formation. The first decision serves to generate outer cells, which will differentiate into the trophectoderm; and the inner cell mass (ICM), which will then undertake the second cell fate decision to give rise to epiblast and primitive endoderm (Fig. 1a) ^16,17^.

**Fig. 1:**
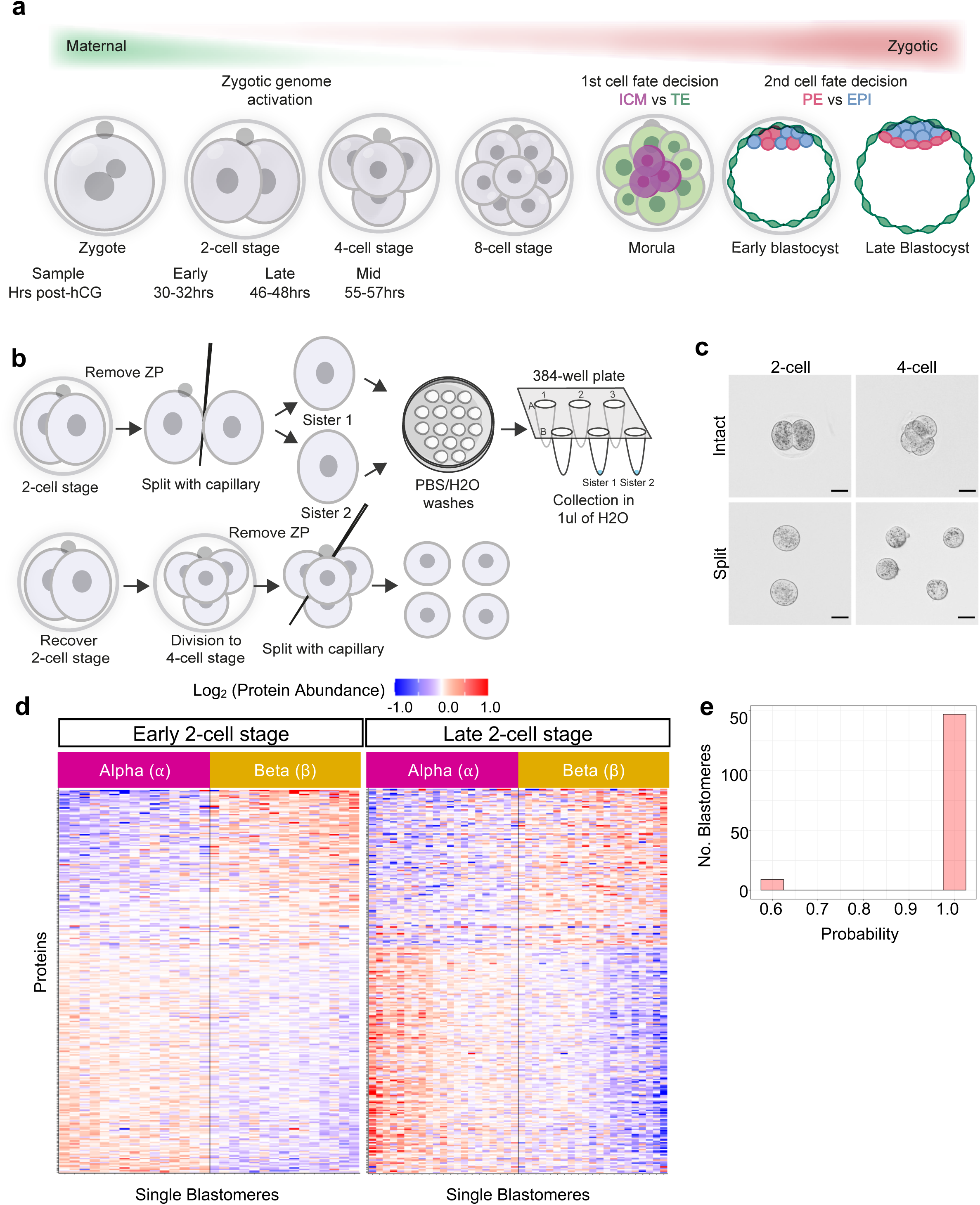
Proteomic asymmetry at the 2- and 4-cell stage mouse embryos. **a,** A schematic of pre-implantation development. Following a series of cleavage divisions, the embryo polarizes at the 8-cell stage and undergoes a series of asymmetric and symmetric divisions to give rise to the inner cell mass (ICM, purple) and outer trophectoderm (TE, green) cells. The ICM then gives rise to the epiblast (EPI, blue and primitive endoderm (PE, pink). Samples were collected at the indicated timepoints. hCG, human chorionic gonadotrophin. **b,** A schematic showing the experimental harvesting of single blastomeres from 2-cell (top) and 4-cell stage (bottom) embryos for single-cell proteomics analysis. ZP, zona pellucida. **c,** Representative images of embryos prior to and following splitting into individual blastomeres. Scale bars, 40 μm. **d,** K-means clustering of 2-cell stage blastomeres results in a consistent bi-clustering of sister blastomeres, i.e., sisters from the same embryo fall into opposing clusters, which we term alpha and beta. Heatmaps of ∼300 proteins with differential abundance in alpha and beta cells. **e,** By changing the starting centroids in the k-means clustering approach 200 times, we obtain vectors of cell cluster classification for each iteration. This allows us to determine the probability of cells landing in the same cluster (alpha or beta). The majority of embryos consistently fall into the same cluster, indicating that the clustering approach is stable.

For decades, it was thought that all blastomeres had the same developmental potential to form these three lineages until the 16-cell stage^5,6,18^. However, this view has changed over time, with multiple lines of evidence suggesting that totipotency is not only lost gradually, but also unevenly, in blastomeres before lineage specification^10, 12, 15,19–22^. First, lineage tracing indicates that blastomeres of 2-cell mouse contribute unevenly to the ICM versus the trophectoderm in the blastocyst^8,9,12^. Second, splitting of 2-cell mouse embryos into monozygotic twin “half embryos” found that one blastomere retains totipotency and gives rise to a live mouse, while the other cell fails to do so in the majority of cases^7,10^. Lineage analyses of somatic mutations across human tissues with different developmental origins, including the placenta, suggest that such uneven contribution arising from the 2-cell blastomeres may also exist in humans^22–26^. Indeed, the most recent lineage tracing studies have shown that, in most human embryos, the majority of the epiblast is derived from only one blastomere of the 2-cell embryo whereas the placenta is derived from both 2-cell stage blastomeres^12^. However, the molecular basis for this asymmetry remains a long-standing question.

In other models, such as *Drosophila melanogaster*, symmetry-breaking mechanisms in the embryo involve asymmetric distribution of mRNAs and, in turn, the encoded proteins^28–31^. Single-cell RNA sequencing methods have identified transcripts^32–34^ such as the non-coding RNA LincGET^35^ and Sox21 mRNA^21^, to be differentially abundant in blastomeres of 2-cell and 4-cell stage mouse embryos respectively. However asymmetries in the abundance of specific mRNAs between sister blastomeres may not be consistent^36^ and RNA abundance does not necessarily reflect protein abundance, as seen across tissues^37^ and during development^38^.

To what extent the proteome differs between individual mammalian blastomeres remains unknown. Single-cell mass-spectrometry (MS) previously revealed proteomic differences between blastomeres of the *Xenopus laevis* embryo^39,40^ and among human oocytes^41^. Bulk samples have been utilized to assess changes in the proteome during mouse embryo development^42–44^ but such bulk samples could not be used to discern intra-embryo heterogeneity. Here, we investigated proteomic differences between single blastomeres from mouse and human embryos and their functional role. Remarkably, we discovered early symmetry breaking of the proteome in the mouse and human 2-cell embryo and even within the zygote. Furthermore, we found that these proteome asymmetries predict the developmental potential of blastomeres.

### Proteomic asymmetry in blastomeres of 2-cell and 4-cell embryos

To initiate our study, we aimed to analyze 1) early 2-cell embryos, 2) late 2-cell embryos, when the major wave of zygotic genome activation occurs^45^, and 3) 4-cell embryos (Fig. 1a-c). To this end, we first established a method to separate and serially wash each blastomere from a single embryo in a way that would avoid any protein contaminant from the culture media while retaining the relationship between blastomeres from the same embryo (Fig. 1b, c). To quantify proteins in single blastomeres we used multiplexed Single Cell ProtEomics MS (SCoPE2)^13,46^. In choosing cells for the isobaric carrier material, we first considered using bulk blastomeres; however, this proved to be unfeasible, as each SCoPE2 set would require hundreds of blastomeres to be collected. We reasoned that mouse embryonic stem cells (ESCs) would serve as adequate carrier cells, as carriers can differ from the single cell samples^47^, ESCs represent a derivative of ICM and we were able to harvest many cells at a time.

To determine the relationship between individual 2-cell stage blastomeres, we performed k-means clustering of the protein abundance normalized to the mean of individual 2-cell embryos. Our data were best explained by two clusters, which we termed alpha and beta (Extended Data Fig. 1a). From our analyses of 36 (15 early and 21 late) 2-cell mouse embryos, sister blastomeres were consistently classified into opposing clusters, with each embryo having an alpha and a beta blastomere with high confidence (Fig. 1d and Fig. 1e).

We uncovered a set of 349 proteins that systematically differed in abundance between alpha and beta blastomeres in 2-cell embryos, out of an average of 1043 proteins quantified per single mouse blastomere (Fig. 1d, Extended Data Table 1). These proteins included maternal factors implicated in zygotic genome activation, such as the cortical granule protein Padi6^48–50^ and the ubiquitin E3 ligase RNF114^51,52^, and cytoskeletal regulators such as Rdx^53^ and Cdc42^54,55^, which are involved in the trophectoderm lineage later in development. To our knowledge, this is the first report of systematic differences between the proteomes of sister blastomeres in the 2-cell mouse embryo.

We found that the proteomic differences between sister blastomeres were more pronounced in late 2-cell embryos compared to early 2-cell embryos, as illustrated by the color scale in the heatmap (Fig. 1d). However, the magnitudes of differential protein abundance between sister blastomeres varied (Fig. 1d). We term this varying degree of difference between alpha and beta blastomeres as the degree of asymmetry, which is observed to increase from the early to the late 2-cell stage. Overall quantitation variability of peptides mapping to the same proteins in each blastomere was low and unrelated to the degree of asymmetry (Extended Data Fig. 1b), and thus we infer that this degree of asymmetry is of biological origin.

Next, we investigated 4-cell embryos, whose blastomeres are known to have different molecular and developmental properties^19–21,56–59^. We analyzed 21 4-cell embryos, in which we observed a spectrum of the degree of asymmetry among sister blastomeres from the same embryo (Extended Data Fig. 1c). Previously, we had defined proteins that consistently discerned alpha and beta blastomeres in 2-cell stage embryos: proteins enriched in alpha blastomeres can be called alpha proteins and proteins enriched in beta blastomeres can be called beta proteins. Thus, we can use the quantitation of these particular proteins to quantify the observed differences among sisters. We calculated the variance of the distribution of the alpha and beta proteins in each blastomere, to observe the overall level of variability of protein levels (Extended Data Fig. 1d). In some 4-cell embryos, the level of variance was similar amongst sisters (e.g., embryo wAP539_29), while in others, the levels of variance were more different (e.g., embryo wAP563_24). This reflects the strength of alpha-beta polarization in each embryo. We also calculated the ratio of the mean abundance of alpha and beta proteins in each blastomere and used this ratio to indicate the “strength” of alpha and beta polarisation. This approach indicated a range of strengths per blastomere (Extended Data Fig. 1e). Observing a higher abundance of alpha proteins in a particular blastomere indicates that the blastomere is more alpha. As an example, embryo wAP439_6 has two relatively strong alpha blastomeres and two relatively strong beta blastomeres. In another example, embryo wAP440_7 has one strong alpha blastomeres, a weak beta blastomere, and two strong beta blastomeres.

The 2-cell embryo can generate the 4-cell embryo via four distinct cleavage patterns, defined by the orientation of cell division (meridional – along the animal-vegetal axis or equatorial – perpendicular to animal-vegetal axis, with the animal-vegetal axis defined by the attached second polar body, which is the product of the second meiotic division of the oocyte upon fertilization) and order of division (Extended Data Fig. 2a). The cleavage pattern has been shown to impact the expression of heterogeneous factors at the 4-cell stage^20^ as well as the success of embryo development^19,56^. To investigate whether the alpha-beta composition of the 4-cell embryos was related to a particular cleavage pattern, we labeled one sister blastomere in live 2-cell embryos by micro-injecting synthetic mCherry mRNA, and recorded cleavage division patterns by time-lapse imaging, as described previously^56,60^, collected individual 4-cell blastomeres^56,59^ and performed SCoPE2 (Extended Data Fig. 2b). We found that few proteins and GO terms differed in abundance between alpha and beta blastomere clusters according to cleavage pattern; the statistical power of this analysis was however insufficient to reliably establish these differences (Extended Data Fig. 2c-e).

Overall, we discovered consistent proteomic heterogeneity in sister blastomeres of 2-cell and 4-cell embryos, forming a molecular signature for alpha and beta cell clusters.

### Proteomic asymmetry in the zygote

Having established the alpha-beta asymmetry at the 2- and 4-cell stages, we wondered whether there is asymmetric protein distribution already within the zygote. To test this hypothesis, we manually bisected zygotes meridionally along the animal-vegetal axis, as this is the most frequent orientation of cleavage of the mouse zygote^60–62^. Although the future cleavage plane will not always be recapitulated by experimental bisection of zygotes along the meridional axis, owing to the rotational symmetry of a spherical cell such as the zygote, there should nevertheless be some instances in which the physical cut approximates to the future cleavage plane (Fig. 2a). Thus, we expected that, if alpha-beta differences are already emerging at the zygote stage, some zygote halves will exhibit the alpha-beta protein differences while others will not. Pairs of zygote halves were collected and subsequently prepared and analyzed using SCoPE2 methods (Fig. 2a, b).

**Fig. 2:**
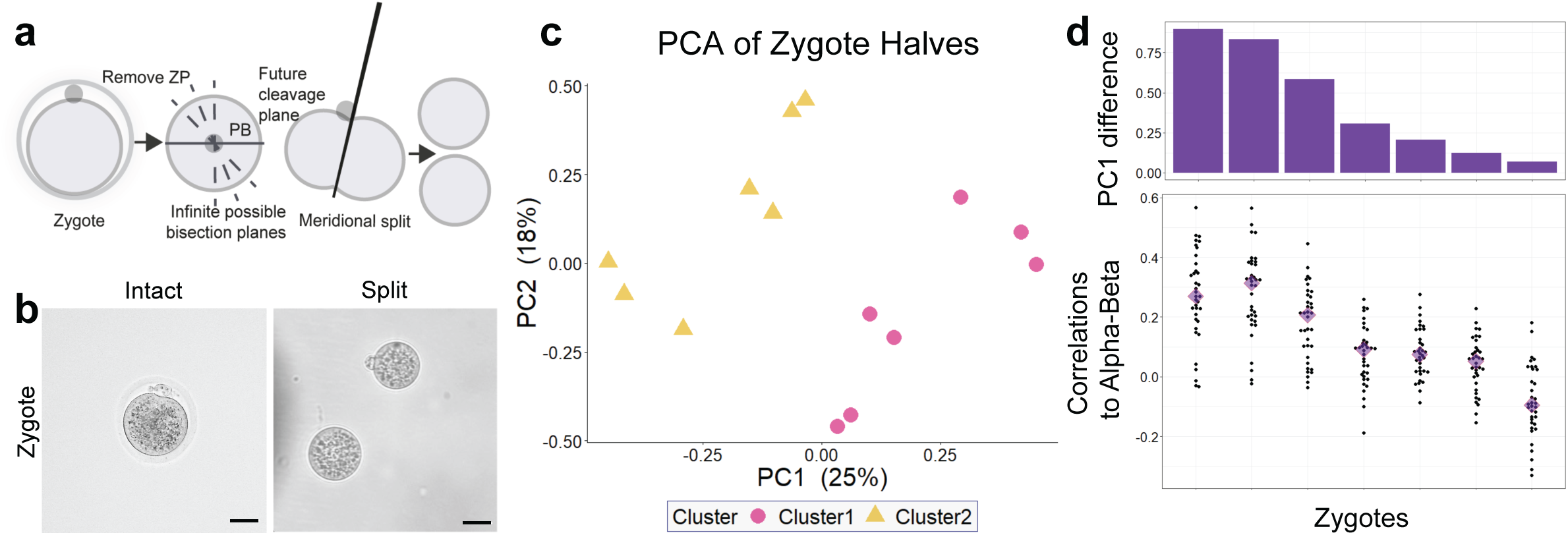
Proteomic asymmetry is inherited from the zygote. **a,** Schematic illustrating the collection of zygotes and subsequent meridional cutting according to the animal-vegetal axis as defined by the position of the second polar body (PB). ZP, zona pellucida. **b,** Representative images of zygotes prior to and following splitting into individual halves. Scale bars, 40 μm. **c,** Principal Component Analysis (PCA) of the zygote halves shows a biclustering pattern. Each zygote pair lands in separate clusters. **d,** The bars on the top show the difference between the PC1 loadings corresponding to each zygote pair ordered in a descending order. On bottom, pairwise spearman correlations were computed between each zygote pair and the 2-cell stage embryos. The correlations were computed on vectors of fold changes of proteins that were both significantly differential between alpha and beta cells and quantified in the zygote dataset. Median correlations of each distribution are shown by triangles.

By performing the same proteomic and clustering analysis as we had done previously for 2-cell stage blastomeres, we found that the zygote halves establish two clusters (Fig. 2c). To test if the different zygote halves were related to the alpha and beta blastomeres, we examined the 172 proteins that were 1) quantified in the zygote halves and 2) had significantly different abundance in alpha and beta blastomeres at the 2-cell stage (Extended Data Table 1). For each protein, we determined the median fold change in alpha versus beta cells and in zygote half 1 versus half 2. We found that the Spearman correlation (r = 0.45) is significant (p-value < 1e-8, Extended Data Fig. 3a). Furthermore, when we took all pairwise correlations of these protein fold-changes between each zygote and embryo, we found that most zygotes had a median positive correlation with magnitude directly proportional to the degree of separation along PC1 (Fig. 2d). This result is consistent with the expectation that variation in the plane of physical cutting along the meridional axis of the zygote will influence the sampled cross-section of protein distributions and thus the magnitude of the correlations. Differences in the proteomes of zygote halves correlate significantly with the protein differences between alpha and beta blastomeres at the 2- and 4-cell stage, which suggests that asymmetry likely stems from differential protein localization in the zygote. Our data point towards the inheritance of asymmetry from the zygote to the 2-cell stage.

As we observe asymmetry within the zygote and during stages prior to or during the major wave of zygotic genome activation, we hypothesize that proteome asymmetry might be driven by post-transcriptional mechanisms concerning maternal transcripts and proteins from the oocyte. When assessing previously published mouse single cell transcriptional data that span the early, mid and late 2 cell stage^63^, we observed stage dependent enrichment of the transcripts associated with alpha and beta proteins (Extended Data Fig. 3b-d). We did not observe a clear transcriptional signature where alpha and beta-associated transcripts show opposing expression patterns between sister blastomeres, but rather stage-dependent expression patterns, potentially reflecting zygotic genome activation (Extended Data Fig. 3e-g). Similarly, pathways which showed differences at the protein level do not exhibit similar patterns when only examining transcripts at this stage (Extended Data Fig. 3h). This suggests post-transcriptional mechanisms and maternal contributions may underlie the proteomic asymmetry we report here.

### Alpha and beta cells are enriched for different biological processes

To decipher processes that could be impacted by the differentially abundance of proteins between alpha and beta blastomeres, we performed protein set enrichment analysis (PSEA). We found that protein degradation and protein transport processes were differentially abundant (q-values < 0.05) in alpha and beta blastomeres (Fig. 3a, b). In particular, ubiquitin- and autophagy-related terms were enriched in beta blastomeres, whereas proteasome-related terms were more enriched in alpha blastomeres. Protein transport terms (channel and signaling-related, molecular motors, and vesicle transport) were enriched in beta blastomeres, with processes related to molecular motors exhibiting the highest median fold difference.

**Fig. 3:**
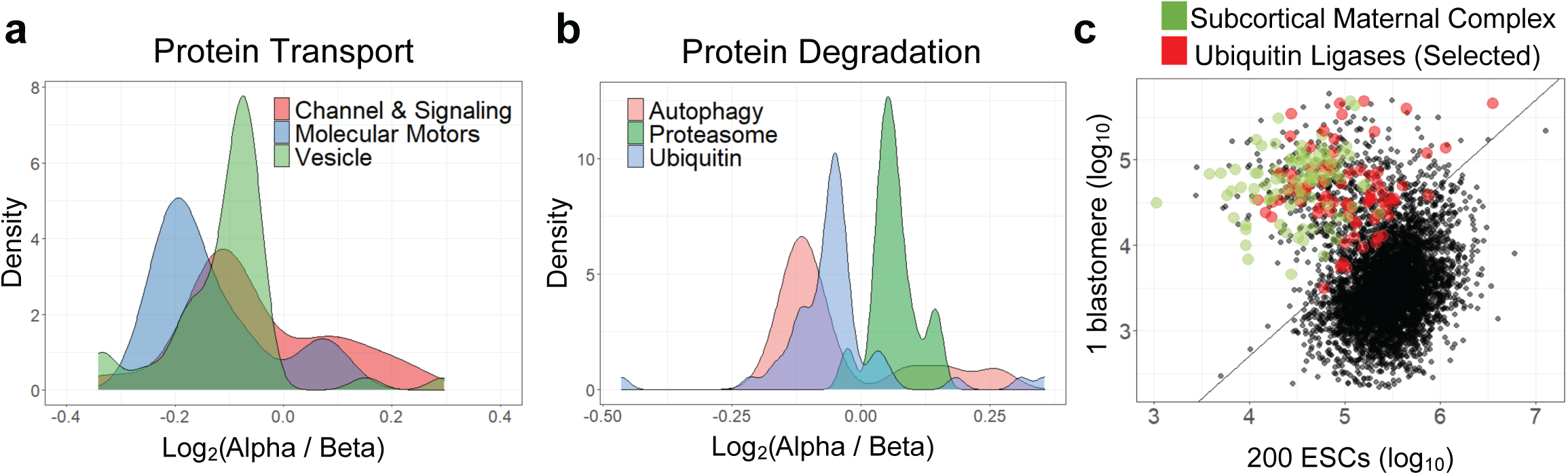
Alpha and beta blastomere clusters exhibit differential biological processes. **a, b,** PSEA analysis revealed differential abundance of proteins related to specific biological processes between alpha and beta cell clusters, namely protein degradation and protein transport. **c,** Representative scatter plot of raw reporter ion intensities from one representative blastomere versus 200 ESCs on the log10 scale. Green points correspond to peptides of proteins mapping to the subcortical maternal complex. Red points correspond to peptides of proteins mapping to different ubiquitin ligases. The diagonal line represents a separation between the two clusters. Such scatterplots were observed for blastomeres across the stages. Upon systematic analysis of all blastomeres in all stages, we found proteins involved in protein degradation and transport to be heavily enriched in blastomeres as compared to mouse ESCs.

To understand how ESC proteomes compare with the blastomere proteomes, we compared the peptide-level data of the carrier and single blastomeres. Upon plotting the levels of shared peptides between single blastomeres and samples of 200 ESCs, we found a cloud of peptides that were much more abundant - up to 10-fold higher - in single blastomeres (a representative plot is shown in Fig. 3c). The overall range of peptide abundances should scale with sample size, and so it was surprising to see many peptides exhibiting much higher abundance in single blastomeres as opposed to 200 ESCs. These peptides derive in part from the subcortical maternal complex (SCMC), a maternally encoded multiprotein complex that is critical for early development^64^. The cloud of outliers also includes peptides related to ubiquitin ligases. This high abundance of ubiquitin ligases is further confirmed by systematic GO term analysis across all single blastomeres, which revealed strong enrichment (relative to ESCs) for peptides implicated in protein degradation and protein transport (Extended Data Fig. 4a), consistent with the known importance of proteasomal degradation during this period of embryonic development, encompassing maternal protein degradation, alongside zygotic genome activation and subsequent novel zygotic protein synthesis^51, 52, 65–67^. As proteins mapping to these similar processes were also found to be differentially abundant between alpha and beta blastomeres, these analyses furthermore underscore their potential association with inter-blastomere proteomic heterogeneity.

### Dynamics of alpha and beta differences

Mouse embryos at the zygote and early 2-cell stage depend largely on maternally inherited cellular components, including proteins, mRNAs and ribosomes, prior to the major wave of zygotic genome activation. Different ribosomal stoichiometries have been suggested to contribute to ribosome-mediated translational control during early embryogenesis^68,69^ and in ESCs^70,71^. Therefore, we assessed whether the ribosomal protein (RP) levels in our samples might be consistent with this hypothesis.

We noticed that the levels of most RPs were slightly, yet statistically significantly, elevated in alpha blastomeres compared to beta cells blastomeres in early 2-cell, late 2-cell and 4-cell embryos (Extended Data Fig. 4b). An exception was RPS27A, whose enrichment in alpha blastomeres increased during development. Proteins involved in translation initiation factors were also more abundant in alpha blastomeres as compared to beta blastomeres, whereas GO terms related to endoplasmic reticulum showed the opposite trend (Extended Data Fig. 4c).

To explore whether differences between alpha and beta blastomeres change during development, we first calculated the Euclidean distances between alpha and beta blastomeres from the same embryo, using proteins that were quantified in every cell analyzed. As noted in Fig. 1d, we observed that the degree of proteomic differences between alpha and beta cell clusters increased significantly during early development (Extended Data Fig. 4d), suggesting sisters may increasingly diverge across stages.

The increased sister divergence across time can be attributed to proteins that are either consistently decreasing or increasing across the developmental stages. We considered the 324 proteins which were quantified in both early and late 2-cell embryos and also deemed to be differentially abundant between alpha and beta blastomeres, and found that 278 proteins preserve cluster identity and are either consistently increasing or decreasing over time. When we also take into account the 4-cell stage, a smaller number of proteins (254) continue to preserve cluster identity, of which 108 are monotonically changing across the three timepoints.

To explore which processes are showing monotonic changes across time, we performed PSEA using Spearman correlations across the three developmental timepoints and the corresponding protein fold-changes between alpha and beta cells from each embryo. Representative protein sets that were decreasing in magnitude across the stages were thioredoxin peroxidase activity and dATP binding (Extended Data Fig. 4e, f). We also find that the proteasome regulatory particle is increasing in magnitude within alpha blastomeres, just as we found in our previous PSEA (Fig. 2b). In addition, representative processes of aspartate metabolic and DNA helicase activity exhibit the same behavior. Overall, these analyses attempted to discern which molecular pathways could be driving the increased divergence between alpha and beta as development progresses.

### The role of beta proteins in lineage fate

We investigated a role for three of the differentially abundant proteins in lineage specification. Specifically, we chose Nedd8, Gps1 and PSMC4 based on their differential abundance between alpha and beta blastomeres (Extended Data Table 2 and 3) and their putative roles in preimplantation development. For instance, activation of the ubiquitin-like protein Nedd8, has been implicated in formation of the ICM^72^. Gps1 (Cops1) is a subunit of the COP9 signalosome^73^, which is involved in deneddylation and has been implicated in naive pluripotency and epiblast survival^74–76^. Gps1 is also a putative regulator of expression of the transcription factor Oct4 in human ESCs^77^, but has not been studied in mammalian embryos before. PSMC4 is a component of the 26S proteasome, and a previous report showed that PSMC4 deficient embryos do not develop into blastocysts^78^. Nedd8 and Gps1 have higher abundance in beta blastomeres, whilst PSMC4 has a higher abundance in alpha blastomeres (Extended Data Table 2).

We used RNAi to knockdown (KD) candidates in one blastomere of the 2-cell embryo and evaluated the consequences for lineage contribution and development of the embryo for 3 days, until the blastocyst stage. Briefly, one of the two blastomeres was randomly micro-injected with dsRNA targeting each candidate (or eGFP as a control) and also with mRNA encoding Gap43-RFP to label and follow the progeny of the injected blastomere, using a protocol we previously established^79,80^ (Fig. 4a). Following micro-injection, embryos were cultured to the late blastocyst stage, fixed and stained for Cdx2 to identify the trophectoderm lineage and Sox17 to identify the primitive endoderm (Fig. 4b, Extended Data Fig. 5a). We validated the KD efficiency by micro-injecting the dsRNA into zygotes, and observing a reduction in mRNA levels 48 hrs. later in 8-cell embryos by qRT-PCR (Extended Data Fig. 6a-d). As a complementary approach, we performed overexpression (OE) studies, in which one blastomere of a 2-cell embryo was co-injected with mRNAs encoding the candidate protein fused to an HA tag and a Gap43-RFP to label the progeny of the injected cell, and cultured to the late blastocyst stage (Fig. 4a and c). The OE of candidates was validated by assessment of HA tag expression by immunofluorescence (Extended Data Fig. 6e-i).

**Fig. 4:**
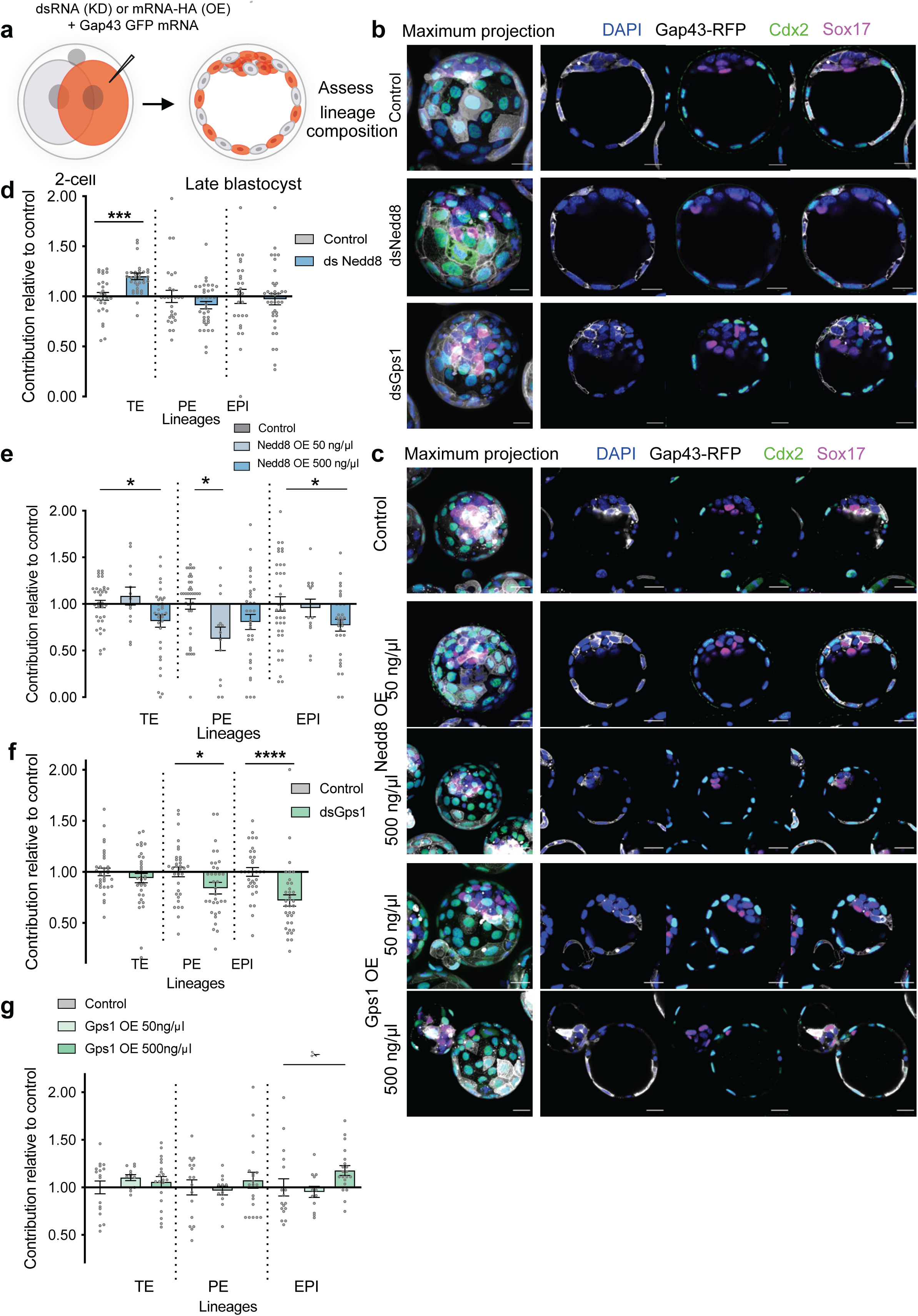
Manipulation of two beta proteins impacts lineage composition. **a,** Schematic of clonal dsRNA-mediated knockdown (KD) or mRNA-mediated overexpression (OE) of candidates. One blastomere of 2-cell stage embryos was injected with dsRNA targeting candidates or eGFP (control) and mRNA for the membrane marker Gap43-RFP for KD experiments. For OE experiments one blastomere of 2-cell stage embryos was injected with mRNA for overexpression of candidates (at the indicated concentration) and for the membrane marker Gap43-RFP. Embryos were cultured to the late blastocyst stage and the contribution of the Gap43-RFP-positive cells to each cell lineage analyzed for both OE and KD experiments. **b,** Representative images of control (ds-eGFP), dsNedd8 and dsGps1 blastocysts. Scale bar, 20 μm. **c,** Representative images of control (Gap43-RFP), Nedd8-HA overexpression (OE) and Gps1-HA OE blastocysts. Scale bar, 20 μm. **d,** dsNedd8 cells show increased contribution to the trophectoderm (TE) lineage. Contribution of dsNedd8 cells to the trophectoderm (TE, Cdx2 positive), primitive endoderm (PE, Sox17 positive), and epiblast (EPI, double negative), was assessed relative to control embryos. Control n = 27 embryos, dsNedd8 n = 36 embryos. Mann-Whitney test, ***p = 0.0005. **e,** Nedd8-HA OE cells show decreased contribution to the TE lineage. Contribution of Nedd8-HA OE cells to the TE, PE and EPI was assessed relative to control embryos. Control n = 36 embryos, Nedd8-HA OE 50 ng/μl n = 11 embryos, Nedd8-HA OE 500 ng/μl n = 28 embryos. Ordinary one-way ANOVA test, adjusted p values, *p = 0.0375 (TE), 0.0120 (PE) and 0.0445 (EPI). **f,** dsGps1 cells show significantly reduced contribution to the EPI. Contribution of dsGps1 cells to the TE, PE and EPI was assessed relative to control embryos. Control n = 33 embryos, dsGps1 n = 35 embryos. Mann-Whitney test, *p = 0.0274, ****p < 0.0001. **g,** Gps1-HA OE cells show significantly increased contribution to the EPI. Contribution of Gps1-HA OE cells to the TE, PE and EPI was assessed relative to control embryos. Control n = 17 embryos, Gps1-HA OE 50 ng/μl n = 12 embryos, Gps1-HA OE 500 ng/μl n = 20 embryos. Kruskal-Wallis test, adjusted p values, *p = 0.0297 . For **d-g**, data are shown as mean ± s.e.m.

We found that the blastocyst cell number and the proportion of RFP-positive cells in the blastocyst was slightly but statistically significantly higher upon Nedd8 knockdown (KD) (Extended Data Fig. 5b and c). Moreover, Nedd8 KD significantly increased the frequency of RFP-positive cells in the trophectoderm relative to controls, but did not have a significant effect on the epiblast or primitive endoderm (Fig. 4d). In comparison we found that the total number of cells in the blastocyst and the proportion of RFP-positive cells in the blastocyst did not differ upon Nedd8 OE (Extended Data Fig. 5i and j), but Nedd8 OE significantly decreased the frequency of RFP-positive cells in the trophectoderm and epiblast (Fig. 4e). We infer that Nedd8 may inhibit the specification and/or proliferation of trophectoderm cells.

Gps1 KD did not reduce blastocyst total cell number (Extended Data Fig. 5d) but reduced the proportion of RFP-positive cells in the blastocyst (Extended Data Fig. 5e), particularly in the epiblast, with a less significant reduction in primitive endoderm and no significant reduction in trophectoderm contribution (Fig. 4f). Gps1 OE did not impact the total number of cells in the blastocysts or the proportion of RFP-positive cells contributing (Extended Data Fig. 5k and l) but led to an increase in contribution to the epiblast, rather than a reduction as observed when knocking down Gps1 expression (Fig. 4g). These data suggest that Gps1 may promote specification and/or proliferation of epiblast cells, and are consistent with the suggested role of Gps1 in promoting pluripotency.

We found that PSMC4 KD reduced the cell number and proportion of RFP-positive cells in all lineages of the blastocyst (Extended Data Fig. 4f, g and h), suggesting that PSMC4 may promote proliferation of uncommitted cells.

Thus, expression of Nedd8 and Gps1, which are more abundant in beta blastomeres, impacts the trophectoderm and epiblast lineage, respectively, in the blastocyst. The phenotypes observed suggest that Gps1 and Nedd8 may play a role in promoting the epiblast fate and suppressing the trophectoderm fate respectively. On the other hand, reduction of PSMC4, which is more abundant in alpha blastomeres, impacted all lineages, perhaps reflecting the importance of the proteasome as the downstream effector of protein degradation. Inhibition of the proteasome has also previously been found to delay DNA replication and cleavage divisions^65^, fitting with our observations. These results indicate that the differential proteins identified by SCoPE2 can impact lineage composition, and point towards the importance of protein degradation pathways during preimplantation development.

### The alpha versus beta identity of blastomeres correlates with differences in developmental potential

We and others previously showed that the developmental potential and subsequent fate of 2-cell stage sister blastomeres are unequal^7–12,19,56^. Specifically, separated sister blastomeres of the 2-cell mouse embryo show discordance in their ability to give rise to a viable embryo and show variation in epiblast size^10^, with one blastomere giving rise to more epiblast cells than its sister (Extended Data Fig. 7a-c). To determine if alpha and beta blastomeres differ in their developmental potential, we separated sisters from 2-cell embryos and analyzed one sister by MS (SCoPE2 or pSCoPE ^81^) to determine its identity (alpha or beta, as quantified by calculating the alpha-beta protein fold change) and cultured the other sister cell to the blastocyst stage (Fig. 5a and b, Extended Data Fig. 7d). We took advantage of our observations that sister blastomeres always fall into opposing clusters, with each 2-cell embryo containing an alpha and a beta cell (Fig. 1d). Thus, we inferred that identity to the cultured sister cell was the opposite to its sister that was analyzed by MS (Fig. 5a). We correlated the identity of the 2-cell blastomere with its subsequent blastocyst development including total cell number, and number of cells in each of the three lineages (Fig. 5c, Extended Data Fig. 7e-h). We found that blastomeres with a higher beta identity gave rise to blastocysts with a higher proportion of epiblast cells (Fig. 5c). Crucially, beta blastomeres are more likely to give rise to blastocysts with 4 epiblast cells, the minimum number required for successful further development^56^ (Fig. 5d). These data agree with our knockdown and overexpression data (Fig. 4), which suggest that beta proteins support epiblast formation and/or inhibit trophectoderm formation. It was recently reported that inheritance of the polar body at the 2-cell stage predicts developmental potential, with the sister inheriting the second polar body, giving rise to more ICM cells^82^. In agreement with this observation, when examining whether there was any relation between inheritance of the polar body and alpha or beta identity, we found that the sister associated with the polar body was significantly more likely to be a beta cell (Extended Data Fig. 7i).

**Fig. 5.**
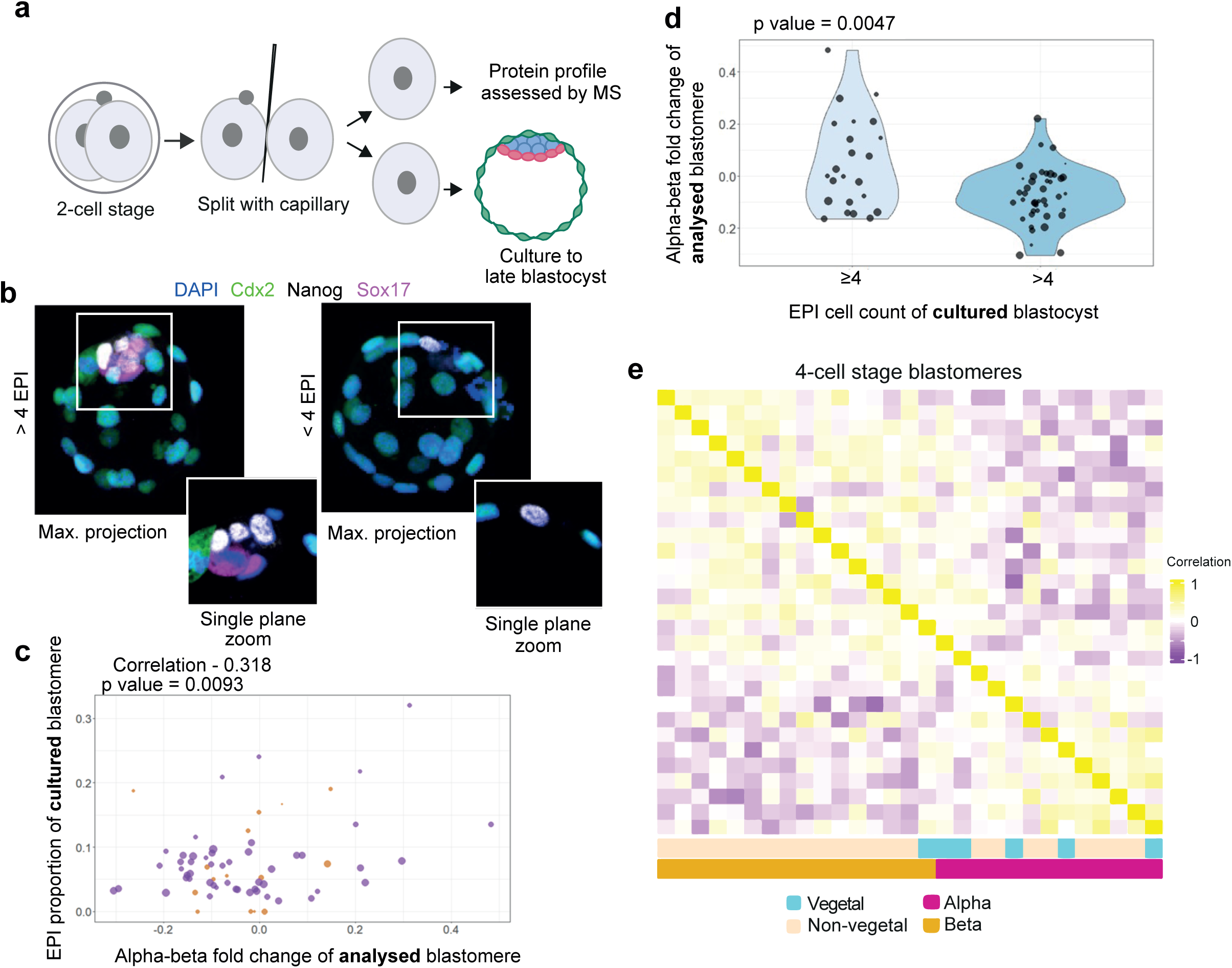
Beta cells have a higher developmental potential. **a,** Schematic illustrating the collection of one sister blastomere for single cell proteomics analysis and subsequent culturing of the other sister to the blastocyst stage. **b,** Representative images of blastocysts with 4 or more epiblast (EPI) cells and fewer than 4 EPI cells. Images are shown as maximum projections and representative single plane zooms showing the composition of the inner cell mass. **c**, Normalized EPI cell counts trend positively with sister cells’ alpha-beta polarization. Plot shows paired blastomere data that was filtered for the sister blastocyst’s characteristics, i.e., at least 10 cells total and have only zero or one lineage totally absent. The size of the data points corresponds to the total number of cells present in the resultant embryo from the blastomere that was left in culture. The color of the data points corresponds to the presence of the three lineages: purple dots mark a blastocyst that contained all three lineages, while brown dots mark a resultant blastocyst that contained at least two lineages. The relationship between the number of epiblast cells in the resulting blastocyst and the corresponding sister’s alpha-beta polarization is quantified by a Pearson correlation computed using all displayed datapoints. **d**, Violin plots of healthy vs less healthy blastocysts and their sister’s alpha-beta polarization. Only blastocysts with more than 10 cells and at most 1 lineage absent were included in this analysis. Healthy blastocysts are defined as having at least 4 epiblast cells, Less healthy as having 3 or fewer epiblast cells. The size of the points is proportional to the total cell count of the resulting blastocyst that was imaged. The statistical significance of this result was tested with a t-test that had a resulting p-value of 0.005. **e.** Heatmap showing the pairwise cell correlations (based on vectors of alpha-beta proteins that exhibited high fold-change), for 4-cell stage blastomeres. Each tile represents a correlation value between two blastomeres, while the color bars below indicate whether the blastomere was identified as a vegetal cell, and its alpha-beta polarization. From this analysis, two clusters of blastomeres can be observed corresponding to alpha and beta. Vegetal cells are significantly more likely to cluster with the alpha-like cells (p=0.047, as calculated using the hypergeometric distribution probability).

At the 4-cell stage, vegetal blastomeres (the blastomere that is furthest away from the polar body, which defines the animal pole of the embryo), are known to be significantly biased to the trophectoderm and have a lower developmental potential^19,56^. We therefore examined the alpha-beta classification of vegetal blastomeres in our 4-cell stage embryos (Fig. 5e). Plotting the pairwise cell correlations for blastomeres from 4-cell embryos, revealed two clusters corresponding to the alpha-beta polarization of each blastomere. We found that vegetal blastomeres were significantly more likely to be alpha than their non-vegetal counterparts. Thus, the alpha-beta identity of a blastomere correlates with differences in developmental potential, with beta blastomeres having a higher developmental potential and alpha blastomeres a lower developmental potential.

Since 2-cell stage blastomeres are often asynchronous in their cell cycle progression, we wondered whether the divergence of alpha and beta blastomeres can be correlated with the asynchrony in developmental timing. In order to test this, zygotes were micro-injected with PCNA-clover mRNA, foci of which indicate S phase progression^83,84^, and following cleavage to the 2-cell stage, assessed by live imaging to see which sister had completed S phase and entered G2 first (Extended Data Fig. 8a, b). After imaging, single 2-cell stage blastomeres were collected for subsequent MS analysis using pSCoPE^81^. When normalizing the MS data within each embryo, sister blastomeres consistently fell into opposite clusters, as observed before. However, we found that the distributions of alpha-beta polarization are not significantly different between ‘early’ and ‘later’ exit from S-phase, and therefore we do not see a relationship between exit from S-phase and alpha-beta identity (Extended Data Fig. 8c).

Concurrently, the experiment was repeated but blastomeres were allowed to develop further to the blastocyst stage, at which point they were fixed and stained for lineage markers (Extended Data Fig. 8d). As alpha and beta blastomeres give rise to blastocysts with differing cell numbers of epiblast cells (Fig. 5c and d), we compared the epiblast cell number from blastocysts arising from sister 2-cell blastomeres which completed S phase ‘first’ or ‘second’. No difference in the distribution of epiblast cell numbers was observed (Extended Data Fig. 8e). This supports our finding that cell cycle asynchrony, as assessed by S phase exit, is unlikely to differentiate alpha and beta cells.

### Proteome asymmetry is conserved in human 2-cell stage embryos

We next investigated if the protein patterns defining alpha and beta blastomeres in mouse embryos are conserved in human embryos. To this end, we examined 2-cell human embryos, which were donated to our research via IVF clinics (Fig. 6a). Since access to human embryos at the early cleavage stages is extremely limited for technical reasons, the number of embryos examined was fewer than for mouse. As these samples are extremely precious, we used two orthogonal single-cell MS methods: label-free data-independent acquisition or SCoPE2 data-dependent acquisition methods. These methods have different systematic biases, and thus concordant results are unlikely to be due to a methodological bias^85^.

**Fig. 6:**
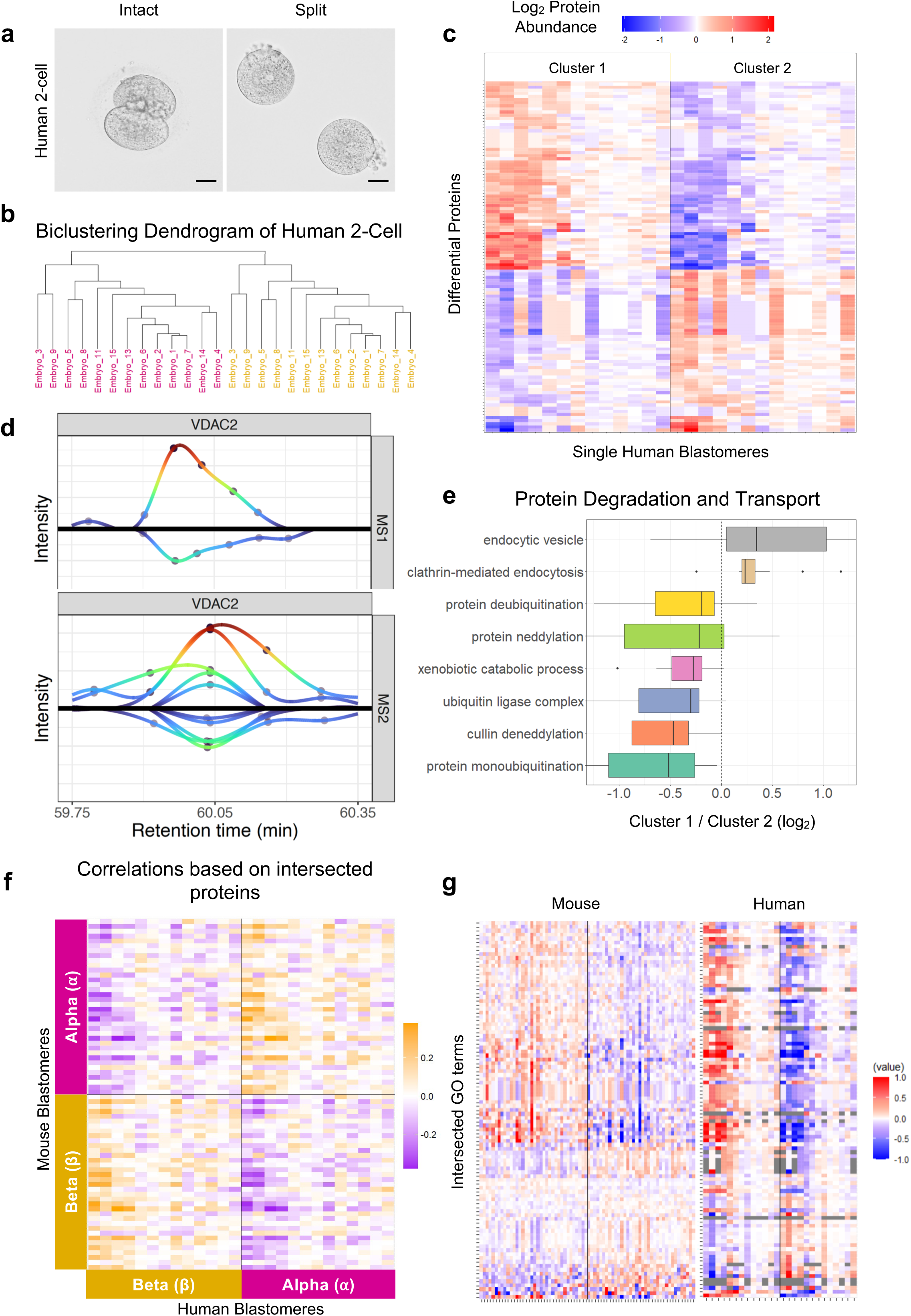
Alpha and beta clusters are conserved in human embryos at the 2-cell stage. **a,** Representative images of human 2-cell embryos prior to and following splitting into individual blastomeres. Scale bars, 40 μm. **b,** Dendrogram illustrating the biclustering behavior in all processed human 2-cell stage blastomeres. Embryo numbers are for indexing purposes only. Color coding indicates the cluster in which each blastomere is classified. **c,** Heatmap of the 113 proteins that are differentially abundant between the two cell clusters. Human blastomeres on the x-axis are ordered in the same way as the dendrogram, while the proteins on the y-axis have been ordered through hierarchical clustering. **c,** Extracted ion chromatogram of peptide mapping to VDAC2 on both the MS1 and MS2 levels indicate consistent fold change between sister cells. **e,** Boxplots of fold changes between sisters of proteins contributing to protein degradation and protein transport terms, which were found to be significantly differential between alpha and beta cells in the human 2-cell stage dataset. **f,** Heatmap of pairwise correlations among mouse and human 2-cell stage embryos based on all intersected proteins shows two clusters, and hence, the level of agreement between alpha and beta classification is positive. **g,** Heatmap of intersected GO terms that are significantly differential between mouse and human (each value represents z-score of median protein abundance for each GO term in each blastomere).

We performed the k-means clustering approach as implemented with the mouse blastomeres and remarkably, we found that sister 2-cell human blastomeres also fell into two opposing clusters (Fig. 6b). Between the two cell clusters, we identified 113 differentially abundant proteins at 1% FDR (Fig. 6c, Extended Data Table 4). To assess the raw MS data more closely, we extracted the ion chromatogram (XIC) for one of the proteins with the highest fold difference, VDAC2, a voltage-dependent anion-selective channel protein. Both the MS1 and the MS2 XICs indicate consistent differential abundance of VDAC2 across the sister blastomeres (Fig. 6d). This observation of VDAC2 being increased in one blastomere fits with our observations from mouse embryos in which protein transport was highly differential between alpha and beta blastomeres. The functional validation of the role of the identified differentially distributed proteins in human embryos at such an early developmental stage is unfeasible.

To further characterize the clusters of human blastomeres, we performed PSEA. Similar to the corresponding analysis of mouse blastomeres, the results again indicated that proteins involved in degradation and transport were differentially abundant in sister blastomeres of 2-cell human embryos (Fig. 6e). Ubiquitin-related proteins were enriched in one cell cluster type, whereas vesicle-related proteins were enriched in the opposing cluster, similar to mouse alpha versus beta blastomeres, respectively.

The two clusters observed in human embryos suggest conservation of the existence of proteomic heterogeneity at the 2-cell stage across human and mouse, therefore we sought to test the concordance more directly. Using 877 proteins whose orthologues were quantified in both mouse and human 2-cell stage embryos, we calculated the pairwise correlations between mouse and human blastomeres (Fig. 6f). The results indicated two distinct clusters, which support that alpha-beta intra-embryo protein differences are conserved across mouse and human and allowed us to extend the alpha and beta annotation to human blastomeres. Additionally, we examined the GO terms that were significantly differential between alpha and beta cell clusters and that were shared between the mouse and human data and found significant concordance among the GO term directionality in the two organisms (Fig. 6g).

To further understand the similarities and differences between the mouse and human blastomeres, we narrowed our space of analysis into proteins that were found significantly differential between alpha and beta 2-cell mouse blastomeres. Then, we estimated the fold changes between the median levels of each protein for alpha and beta cells in both human and mouse. Through the comparison of these fold-changes, we find 68 proteins that change in the same direction in both human and mouse blastomeres, and 98 proteins that exhibit opposite directionality (Extended Data Fig. 9a and b). Such analysis suggests that whilst there is some conservation of how the alpha-beta proteins behave across species, there are also species differences. Therefore, intra-embryo proteomic differences we found in the mouse are also present in human embryos both at the level of differentially abundant proteins and enrichment of protein sets representing different themes of biological processes.

## Discussion

For decades, it was thought that the blastomeres of mouse and human embryos are equivalent to each other in their developmental properties until reaching differential positions within the embryo at the 16-cell stage. However, the advent of new technologies to track individual cells in living embryos to determine their developmental fate and potential, and to examine the patterns of gene expression in single cells, has shown that blastomeres can become different from each other at earlier stages of development^12,19,21,35,56,57,86^. Moreover, only one sister blastomere appears to be truly totipotent in the majority of 2-cell mouse embryos when sister blastomeres are separated from each other^7,10^. A central question revolves around the molecular factors generating this heterogeneity. Here, we explored intra-embryo differences from the zygote to the 4-cell stage, using single-cell MS proteomics for the first time.

Unexpectedly we discovered intra-zygotic and inter-blastomere asymmetry of hundreds of proteins in mouse and human embryos. Specifically, we found that sister blastomeres from 2-cell mouse embryos can be consistently classified into two clusters, which we termed alpha and beta. It is worth noting that the collection of such samples was not trivial. All matched blastomeres from each embryo and individual zygote halves must first be separated from each other and then thoroughly washed in pure water without lysing to obtain “clean” MS data, without contaminating spectra from embryo culture medium and allowing for downstream intra-embryo analyses. The use of isobaric mass tags makes it challenging to estimate the reliability of quantification of each protein in each single cell, especially for proteins represented by a single peptide. This challenge was mitigated in the samples analyzed by DIA and can be further mitigated in future studies using plexDIA^85^. Furthermore, utilising both DDA and DIA, two orthogonal methods, we were able to establish a pattern of asymmetry in the human 2-cell and mouse 4-cell embryos, that recapitulates the alpha-beta asymmetry we observe in mouse.

We found that protein asymmetry is already present in the zygote before zygotic genome activation, which suggests that the symmetry breaking mechanism is at least partially driven by mechanisms other than transcription. Our data reveal that proteins involved in protein degradation and protein transport are highly enriched in blastomeres and differentially abundant across alpha and beta cells. We propose that these processes are involved in symmetry breaking in the embryo, and in divergence in developmental potential prior to lineage diversification.

We observed that beta blastomeres have a higher developmental potential and can give rise to a blastocyst with more epiblast cells than alpha blastomeres. Furthermore, we observed that vegetal 4-cell stage blastomeres, which are known to have a lower developmental potential^19,56^ are more likely to be alpha, linking the proteomic asymmetry we have found between sister cells in the earliest stages of development to eventual developmental fate. Finally, we found that proteomic asymmetry appears to be conserved in human 2-cell embryos and that the alpha and beta classification can be applied to human blastomeres.

Throughout the stages examined, we found a range in the extent of alpha-beta polarization which may reflect the underlying variability in the development of the mammalian embryo, for example, different planes of cleavage divisions from the 1- to 2-cell stage and/or the 2- to 4-cell stage may impact the proteomic asymmetry present in daughter cells. Indeed, it has been shown before that the cleavage pattern in relation to the polar body, does affect the fate of mouse embryo blastomeres^19,20,56^. This variability also fits with previous observations that for the majority of mouse 2-cell embryos only one sister blastomere is totipotent and able to rise to a live mouse, but that there are nonetheless a minority of embryos in which both sisters retain totipotency^7, 10^.

The potential mechanisms underlying this early symmetry breaking during the earliest stages of mammalian development require future investigations. It would be of interest to determine if the asymmetric distribution of proteins in the zygote is maternally or paternally driven, meaning arising already in the oocyte or only following fertilization or both. Intriguingly we find components of cytoplasmic lattices to be asymmetric, with such cytoplasmic lattices being recently implicated in the storage of maternal proteins in the mouse oocyte^87^. Two of the lattice proteins (Padi6 and Ooep) show a biased accumulation in beta cells, and many of the proteins that accumulate on the lattices show a biased distribution in alpha and beta cells. For example, more ribosomal proteins are present in alpha cells whereas mitochondrial proteins, peroxidase activity, tubulin and 14-3-3 proteins accumulate in the beta cells. Overall these findings point to possible role for the maternal transmission of proteins to the embryo, with distinct roles for these maternal proteins in biasing cell fate from the 2-cell stage. Such asymmetric protein distribution may be later paired with asynchronous zygotic genome activation between the blastomeres, leading to transcriptional differences between sisters that may compound the differences we observe from the 2- to 4-cell stage. In summary, our single-cell proteomics approaches revealed the earliest incidence of proteomic asymmetry in the mammalian embryo, which is correlated with developmental potential, providing novel insight into the role of early heterogeneity and cell fate.

## Extended data figure legends

**Extended Data Fig. 1:**
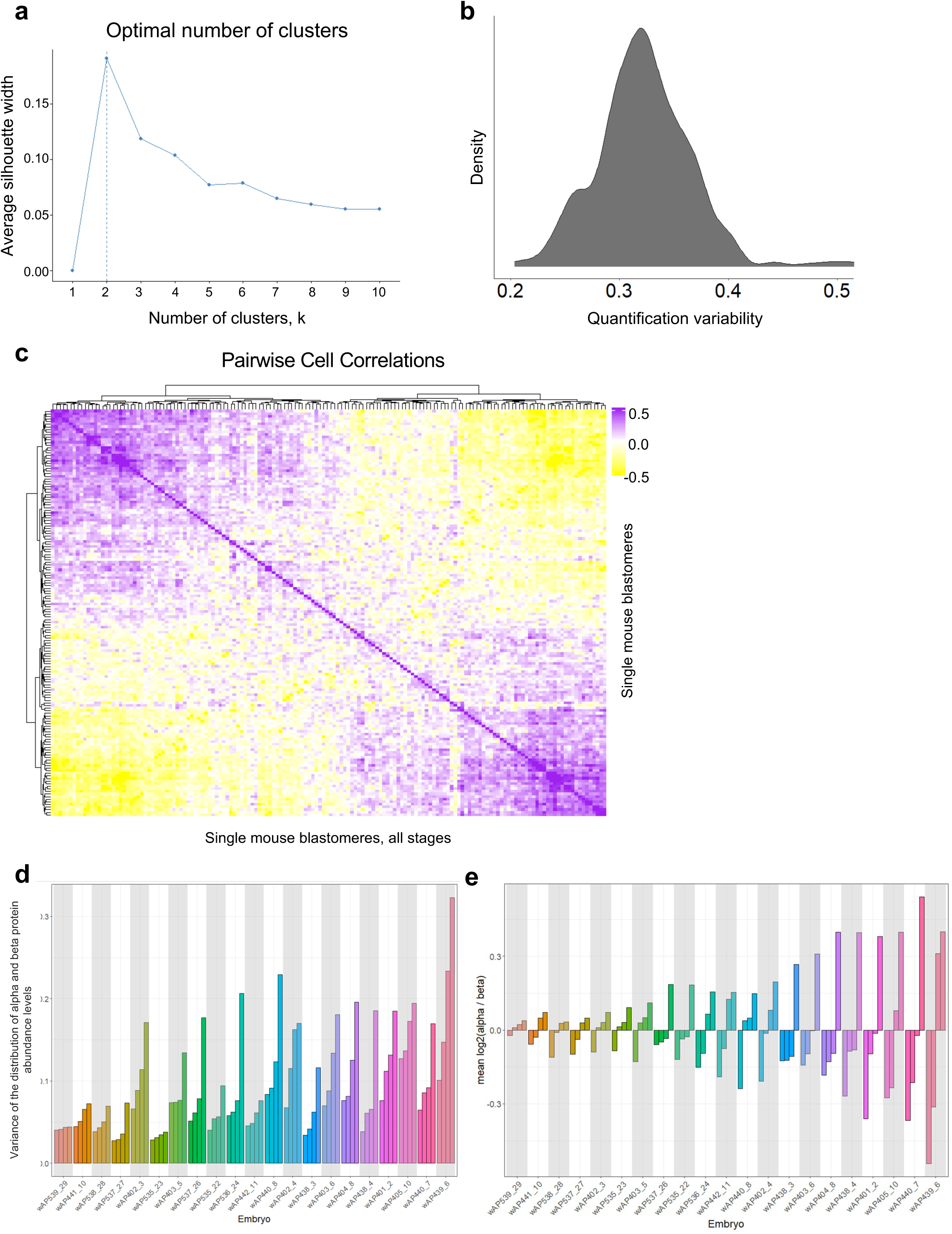
Data Exploration related to Fig. 1. **a,** The number of clusters (k) that can best explain the data plotted against the average silhouette width. In this case, k = 2 provides the best explanation for the data. **b,** Representative density plot showing quantitation variability for peptides mapping to the same protein in each mouse blastomere. **c,** Spearman correlation plot of all individual blastomeres, from the 2-cell and 4-cell stages, which demonstrates the presence of two clusters. **d,** The level of variance of alpha-beta protein quantitation in each blastomere at the 4-cell stage Each grouping of 4 blastomeres represents an embryo on the x-axis. The y-axis is the variance of alpha and beta protein abundances in each blastomere. All blastomeres that were part of the same embryo are colored in the same color. **e,** Variability of alpha-protein quantitation among sisters in each 4-cell stage embryo Each grouping of 4 blastomeres represents an embryo on the x-axis. The y-axis is the fold change between the mean abundances of alpha proteins and beta proteins in each blastomere. All blastomeres that were part of the same embryo are colored in the same color.

**Extended Data Fig. 2:**
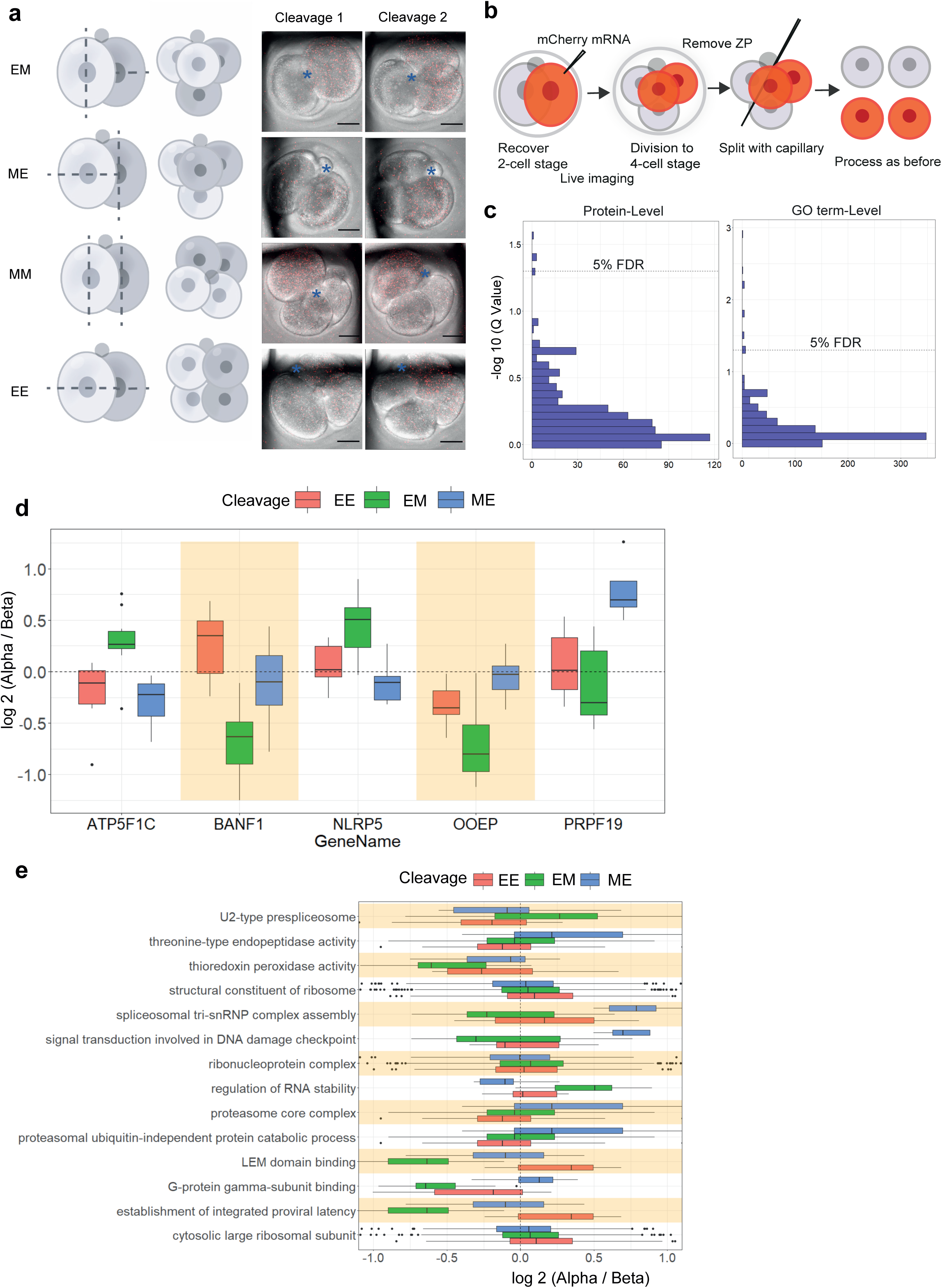
Cleavage Pattern Analysis Between Alpha and Beta cells. **a,** Schematic of division patterns from the 2- to 4-cell stage and representative stills from live imaging to classify division pattern. E denotes equatorial division and M meridional in relation to the animal-vegetal axis of the fertilized egg, with the first letter denoting the first cleavage and the second letter the second. ME (M-division followed by E-division); EM (E-division followed by M-division); MM (consecutive M-divisions); EE (consecutive E-divisions). The position of the polar body is indicated with an asterisk. Scale bar, 20 μm. **b,** Schematic showing the experimental harvesting of single blastomeres from 4-cell stage embryos, with classified division pattern and order, which were subsequently prepared using SCoPE2. Division pattern and order were classified by live imaging, following microinjection of synthetic mCherry mRNA to label one of the two sisters at the 2-cell stage. ZP, zona pellucida. **c,** Q-value distribution of protein-level analysis and GO term-level analysis. Dotted line indicated q.value = 0.05 **d,** Proteins within the 5% FDR threshold in determining differences between alpha and beta cell clusters within the scope of cleavage patterns. **e,** GO terms that passed the 5% FDR threshold in determining differences between alpha and beta cell clusters within the scope of cleavage patterns.

**Extended Data Fig. 3:**
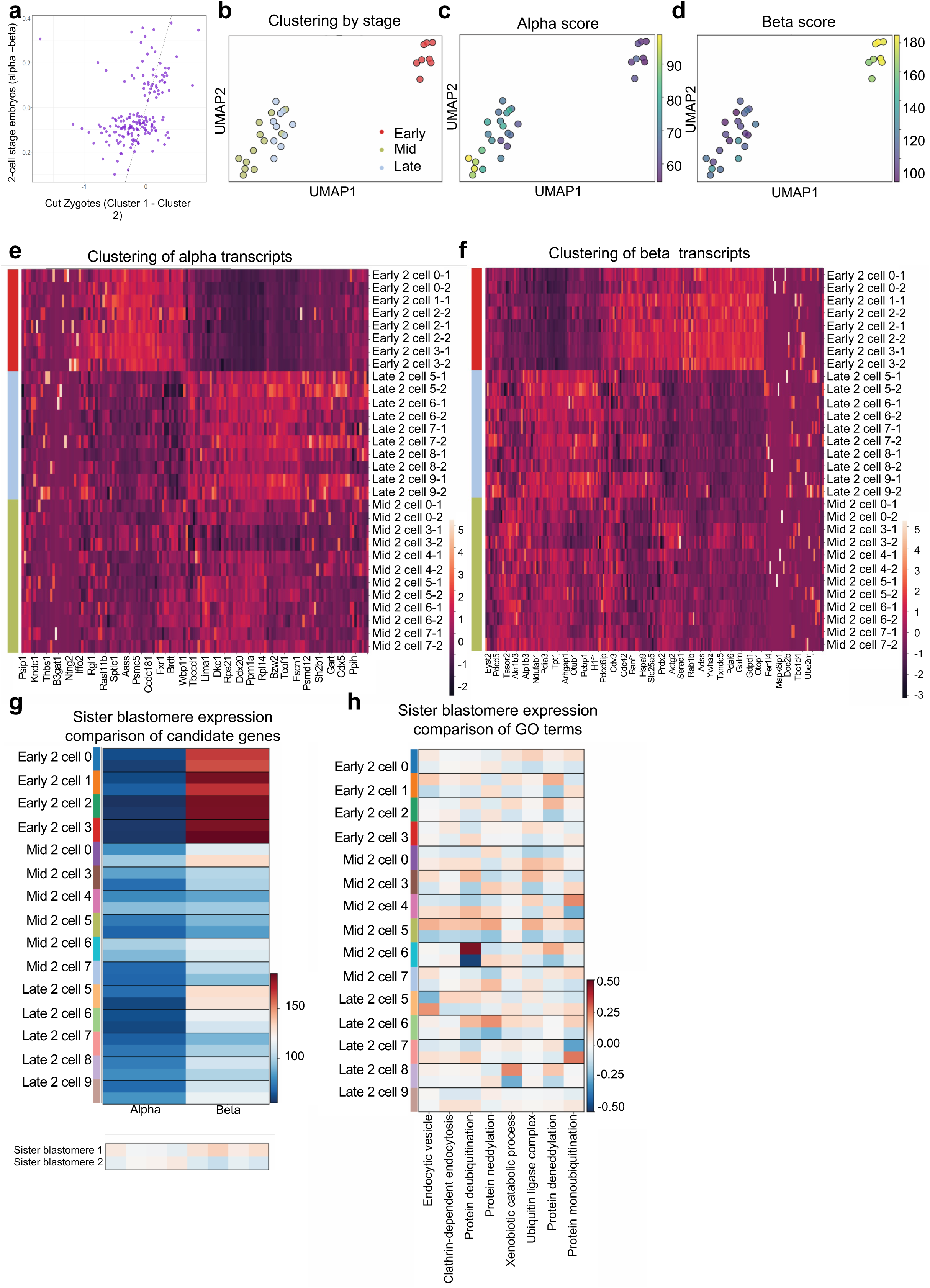
Post-transcriptional mechanisms and maternal contributions may underlie proteomic asymmetries. **a,** Scatterplot of median protein fold changes between zygote halves on the x-axis and median protein fold changes between sister blastomeres at the two-cell stage. The proteins chosen were both differentially abundant between alpha- and beta-type cells at the two-cell stage and quantified in the zygote dataset. We see a positive correlation of -0.45 which is highly significant (p-value < 1e-8). **b,** Uniform manifold approximation and projection (UMAP) of single cell transcriptome of 2-cell embryos from the Deng *et al.* dataset. Individual cells are coloured based upon stage: Early 2-cell in red, mid 2-cell in light green, and late 2-cell in light blue. **c, d,** UMAP portraying either an Alpha (**c**) or Beta (**d**) score for each cell. **e, f,** Clustermap displaying expression levels for Alpha (**e**) or Beta (**f**) transcripts in each blastomere. Sister blastomeres appear next to one another along the vertical axis; blastomeres are labelled by stage, embryo number, and blastomere number, respectively. **g,** Heat map showing Alpha or Beta scores for each blastomere. Blastomeres from the same embryo are grouped together. **h,** Heatmap showing median transcript abundance mapping to particular GO terms that are known to have heterogeneity between blastomeres in the 2- cell embryo based on proteome data. Blastomeres of the same embryo are plotted next to each other.

**Extended Data Fig. 4:**
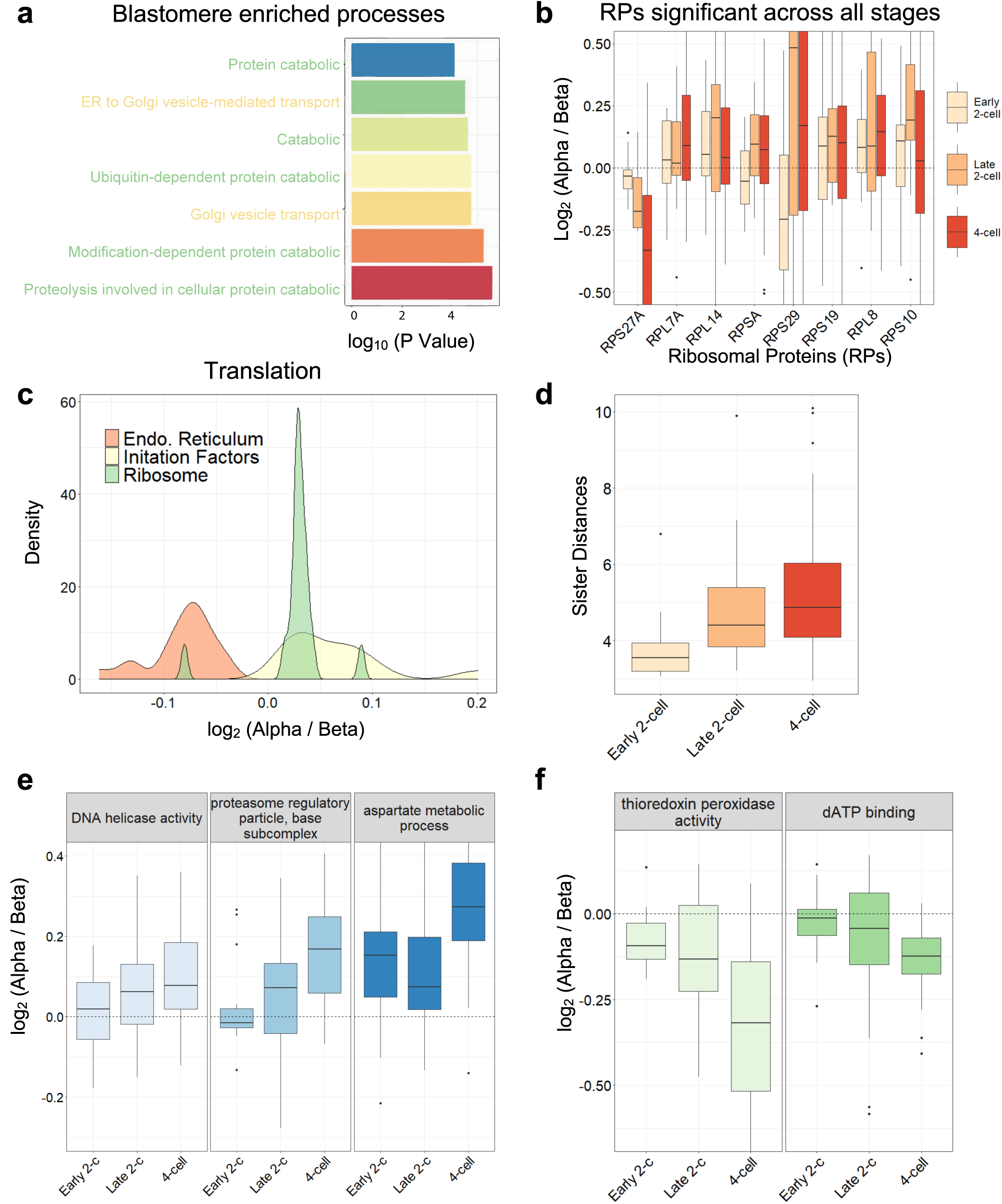
Temporal overview of differences between alpha and beta cells. **a,** P values of the top most enriched processes in blastomeres relative to ESCs. Green font corresponds to protein degradation processes, while yellow font corresponds to protein transport processes. **b,** Boxplots illustrating the levels of significantly differential (1% FDR) Ribosomal Proteins (RPs) between alpha and beta cells. The color of the boxplots corresponds to the developmental stage. RPs were tested separately between alpha and beta cells, and for each stage. **c,** Density plots of protein translation themes that were found significantly differential between alpha and beta cell clusters. **d,** Euclidean distance of normalized protein abundance between each blastomere in each embryo, showing increasing inter-blastomere differences. **e, f,** Correlation values of top protein sets (by highest absolute correlation value) obtained from analysis across the stages.

**Extended Data Fig. 5:**
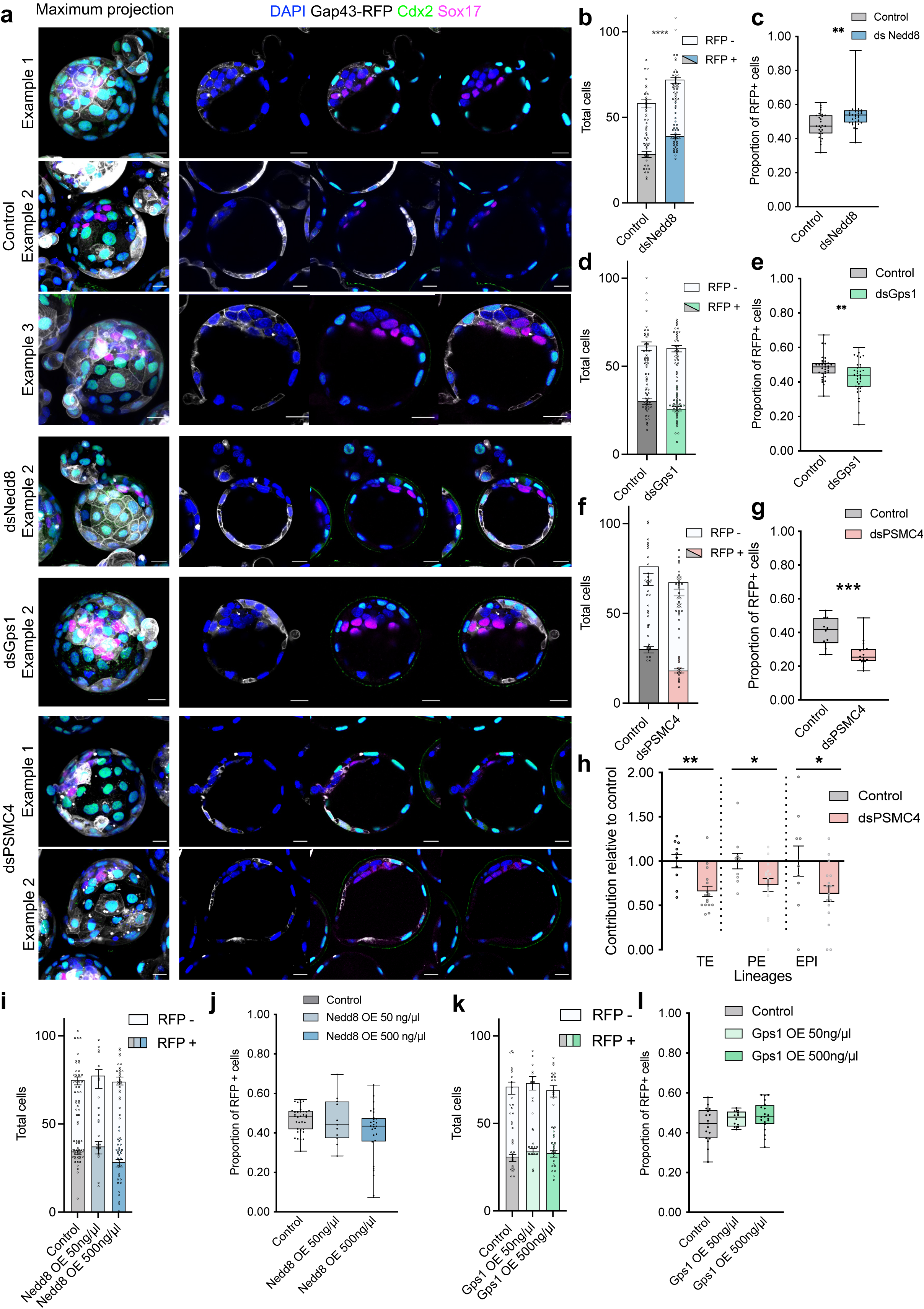
Further data related to Fig. 3. **a,** Representative images further examples of control (ds-eGFP), dsNedd8, dsGps1 and of dsPSMC4 blastocysts. Scale bar, 20 μm. **b,** Bar chart showing the average total number of cells and the proportion of RFP positive or negative cells in control and dsNedd8 late blastocysts. Mann-Whitney test, ****p < 0.0001. **c,** dsNedd8 cells show increased contribution to the blastocyst stage embryo. Control n = 27 embryos, dsNedd8 n = 36 embryos. Mann-Whitney test, *p = 0.0024. **d,** Bar chart showing the average total number of cells and the proportion of RFP positive or negative cells in control and dsGps1 late blastocysts. **e,** dsGps1 cells show significantly reduced contribution to blastocyst stage embryo. Control n = 33 embryos, dsGps1 n = 35 embryos. Mann-Whitney test, **p = 0.0081. **f,** Bar chart showing the average total number of cells and the proportion of RFP positive or negative cells in control and dsPSMC4 late blastocysts. **g,** dsPSMC4 cells show decreased contribution to the blastocyst stage embryo.. Mann-Whitney test, **p < 0.001. **h,** dsPSMC4 cells show decreased contribution to all three lineages. Contribution of dsPSMC4 cells to the trophectoderm (TE, Cdx2 positive), primitive endoderm (PE, Sox17 positive), and epiblast (EPI, double negative), was assessed relative to control embryos. Control n = 10 embryos, dsPSMC4 n = 16 embryos. Mann-Whitney test, *p = 0.0449 (PE), *p = 0.0475 (EPI), **p < 0.0017. Data are shown as mean ± s.e.m. **i,** Bar chart showing the average total number of cells and the proportion of RFP positive or negative cells in control and Nedd8-HA OE late blastocysts. **j,** Nedd8-HA OE cells cells show no difference in contribution to blastocyst stage embryo. Control n = 36 embryos, Nedd8-HA OE 50 ng/μl n = 11 embryos and Nedd8-HA OE 500 ng/μl n = 28 embryos. **k,** Bar chart showing the average total number of cells and the proportion of RFP positive or negative cells in control and Gps1-HA OE late blastocysts. **l,** Gps1-HA OE cells show no difference in contribution to blastocyst stage embryo. Control n = 17 embryos, Gps1-HA OE 50 ng/μl n = 12 embryos, Gps1-HA OE 500 ng/μl n = 20 embryos. The proportion of Gap43-RFP positive cells was assessed in control and mRNA injected embryos in **c, e, g, j** and **l**. For **c, e**, **g, j** and **l**, data are shown as individual data points on a Box and Whiskers plot.

**Extended Data Fig. 6:**
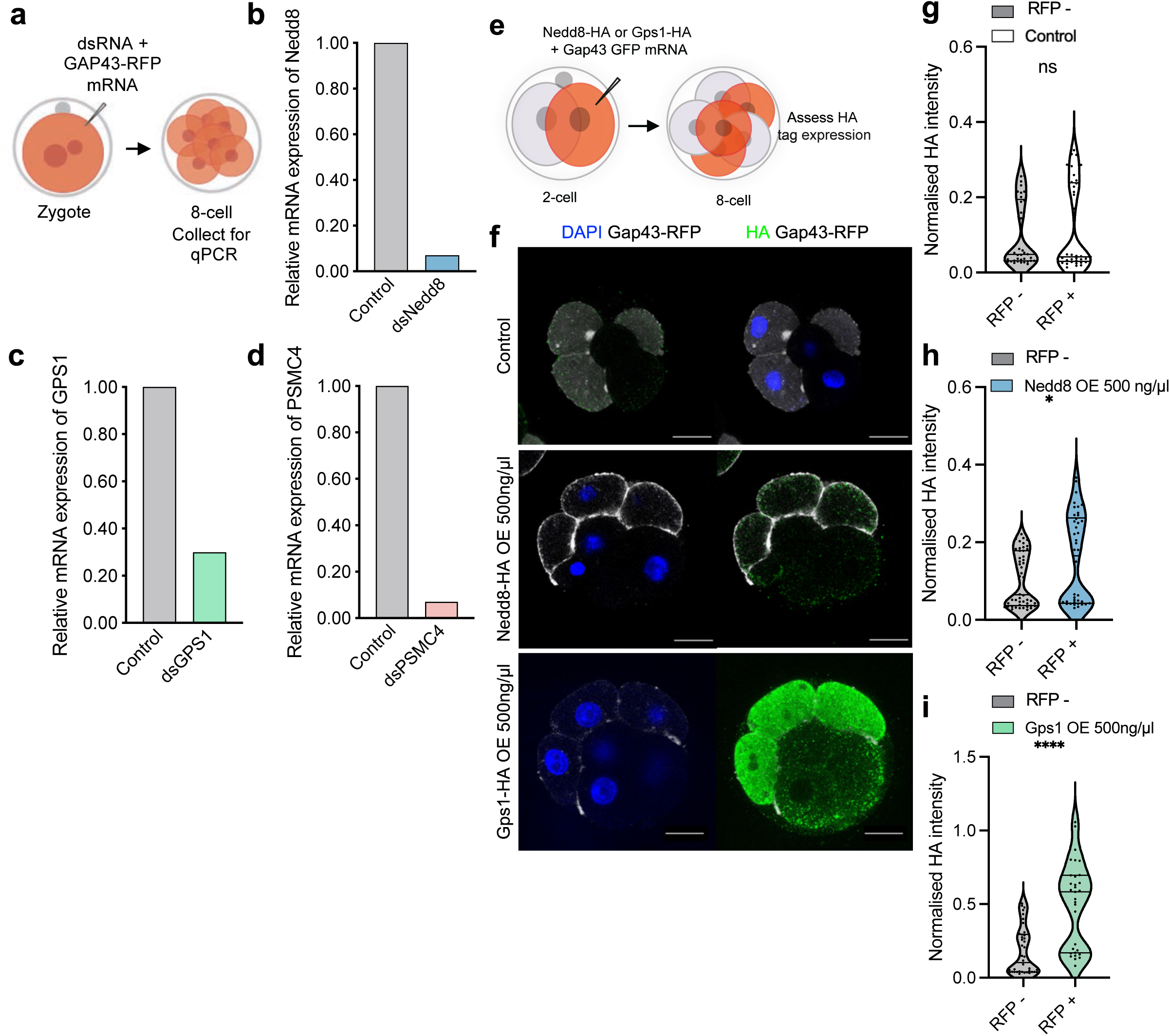
Validation of knockdown and overexpression experiments. **a,** Schematic for validation of dsRNA mediated knockdown. Embryos were injected with dsRNA targeting candidates (Nedd8 = dsNedd8, Gps1 = dsGps1, PSMC4 = dsPSMC4) or eGFP (control) and collected after 48 hrs for qRT-PCR. **b,** Nedd8 mRNA expression was assessed relative to control embryos. Control n = 14 embryos, dsNedd8 n = 14 embryos. **c,** Gps1 mRNA expression was assessed relative to control embryos. Control n = 15 embryos, dsGps1 n = 15 embryos. **d,** PSMC4 mRNA expression was assessed relative to control embryos. Control n = 14 embryos, dsPSMC4 n = 14 embryos. **e,** Schematic for validation of mRNA mediated overexpression. Embryos were injected with mRNA for candidates, tagged with HA (Nedd8 = Nedd8-HA OE, Gps1 = Gps1-HA OE) or Gap43-GFP (control) and cultured for 24 hrs. Embryos were stained for HA and the normalized mean fluorescence intensity values of the Gap43-RFP positive cells and negative cells assessed. **f,** Representative images of control (Gap43-RFP), Nedd8-HA overexpression (OE) and Gps1-HA OE 8-cell embryos. Scale bar, 20 μm. **g, h, i,** HA expression of Gap43-GFP positive and negative cells was assessed in control, Nedd8-HA OE and Gps1-HA OE embryos respectively. Control n = 11 embryos, RFP - n = 40 cells, RFP + n = 37 cells. Nedd8-HA OE n = 12 embryos RFP - n = 48 cells, RFP + n = 42 cells. Gps1-HA OE n = 11 embryos, RFP - n = 42 cells, RFP + n = 31 cells. Student’s t test, *p = 0.0109, ****p < 0.0001. For **g, h** and **i**, data are shown as violin plots.

**Extended Data Fig. 7:**
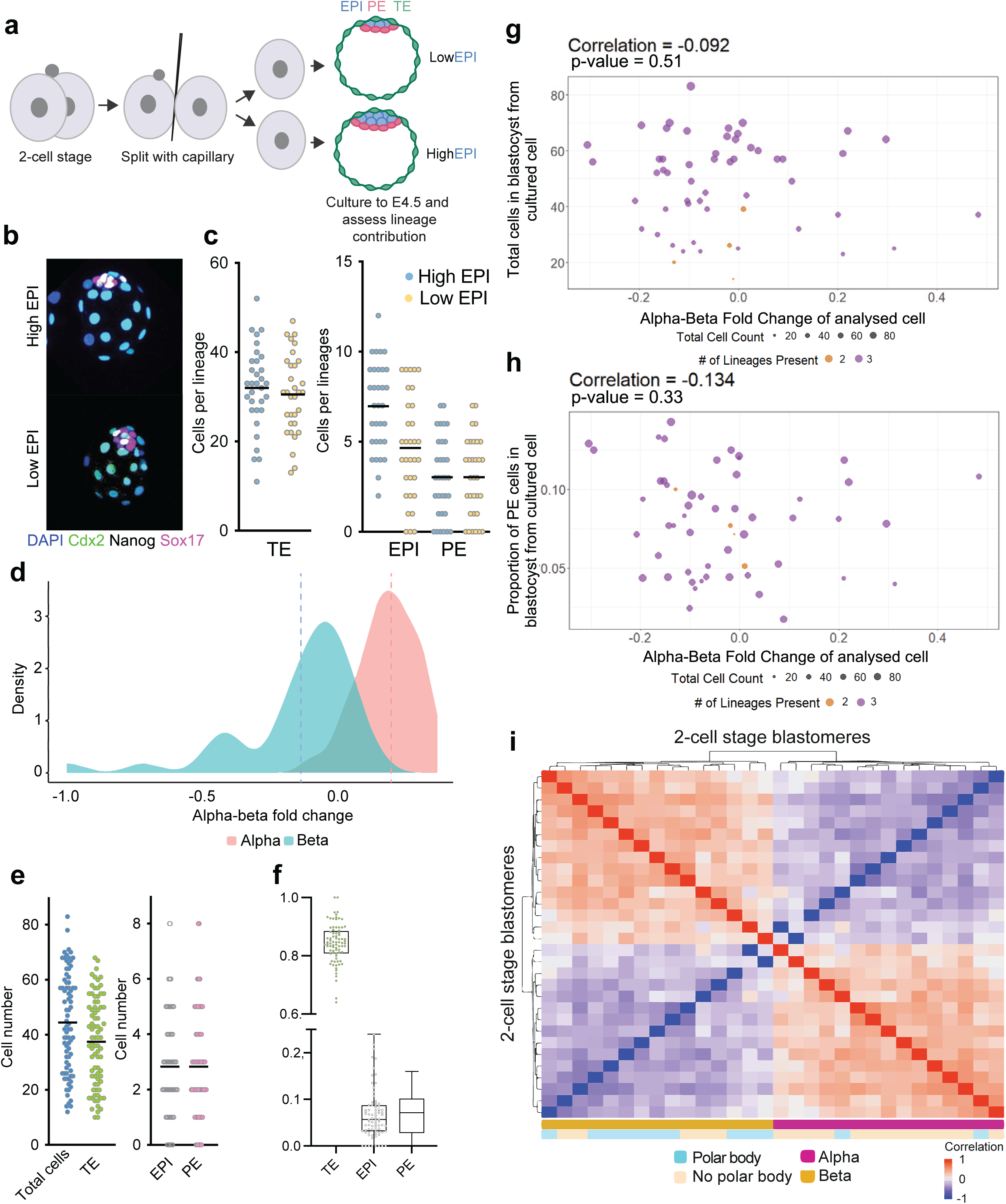
Sister 2-cell blastomeres show differences in their development. **a,** Schematic of split embryo culture. 2-cell embryos were recovered and split before being cultured to the late blastocyst stage in pairs. Blastocysts were assessed for lineage marker expression. **b,** Representative images of a pair of ‘twin’ blastocysts, showing which has more (high) or fewer (low) epiblast (EPI) cells. Images are shown as maximum projections showing the composition of the inner cell mass. **c,** Embryos were classified as high or low EPI within each pair and the number of cells in each lineage (trophectoderm (TE, Cdx2 positive) epiblast (EPI, Nanog positive) and primitive endoderm (PE, SOX17 positive)). n = 32 pairs of blastocysts. **d,** Density plot showing the alpha-beta protein fold changes computed from global normalization. The blastomere type (‘alpha’ or ‘beta’) was defined by K-mean clustering after within embryo normalization for the 2-cell stage samples in which the proteomes of both sisters were analysed. The alpha-beta fold change (x axis) was then computed from blastomere proteomes normalized to the mean across all single 2-cell blastomeres rather within each 2-cell embryo. Colour indicates the clustering based on normalization within each embryo, while the alpha-beta protein fold-change derived from normalizing across all blastomeres is shown on the x-axis. **e, f,** Cell number in each lineage and proportion of each lineage respectively in blastocysts from fig. 4e-h. n = 81 embryos. **g,** Scatterplot of number of total cells in resultant blastocyst versus alpha-beta polarization of sister cell analyzed by MS. Overall, there is no correlation between the two variables. Size of circles indicate the total number of cells in the imaged blastocyst, while the colors indicate the number of lineages present in the imaged blastocysts **h**, Scatterplot of normalized primitive endoderm cell count in resultant blastocyst versus alpha-beta character of sister cell analyzed by MS. Overall, we observe a negative correlation that is not statistically significant. Size of circles indicate the total number of cells in the imaged blastocyst, while the colors indicate the number of lineages present in the imaged blastocysts. **i,** Heatmap showing the pairwise cell correlations for all early 2-cell stage blastomeres. Correlations were calculated based on quantified alpha-beta proteins. Each tile represents a correlation value between two blastomeres, while the color bars below indicate alpha-beta polarization and whether the cell had an associated polar body. Cells with an identified polar body are more likely to cluster with beta cells when including all early 2- cell blastomeres (p=0.0036, as calculated using the hypergeometric distribution probability).

**Extended Data Fig 8:**
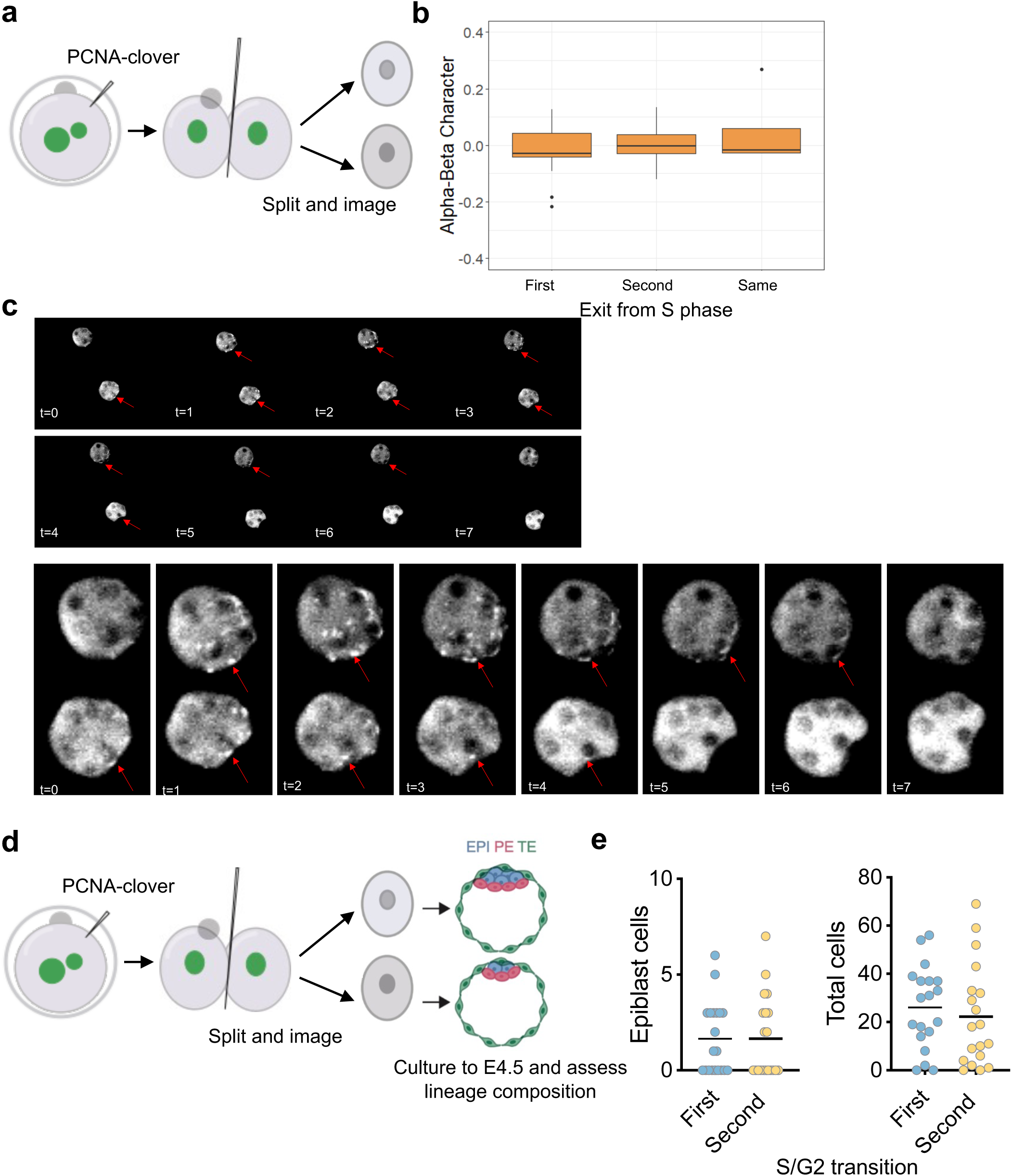
Alpha-beta polarization does not relate to cell cycle asynchrony as assessed by PCNA expression. **a,** Schematic of cell cycle analysis by live imaging of embryos expressing PCNA-clover. Zygotes were injected with PCNA-clover mRNA and allowed to cleave to the 2-cell stage. 2-cell embryos were split and imaged from 36 to 46 hrs post-hCG, during the transition from S to G2 phase, before being collected for single cell proteomics. **b,** Representative still images of live imaging for PCNA-clover expression during the S/G2 transition. Red arrows indicate foci of PCNA-clover, which disappear as the cell enters G2. Magnified views are shown below. Time interval = 15 mins. **c,** Boxplots of alpha-beta polarization (derived from normalization across all single blastomeres) as distributed between blastomeres that exited the S phase either early or late. n = 15 embryos, 30 cells. **d,** Schematic of cell cycle analysis by live imaging of embryos expressing PCNA-clover, followed by culture to the blastocyst stage. Zygotes were injected with PCNA-clover and imaged as in **a**. Following a second cleavage division to the 4-cell stage, split embryos were cultured to the blastocyst stage and lineage composition assessed by immunofluorescence. **e,** Epiblast and total cell number in blastocysts does not show any relation to S/G2 transition. Live imaging was used to determine which sister had ended S phase/entered G2 first or second. n = 38 embryos / 19 pairs.

**Extended Data Fig. 9:**
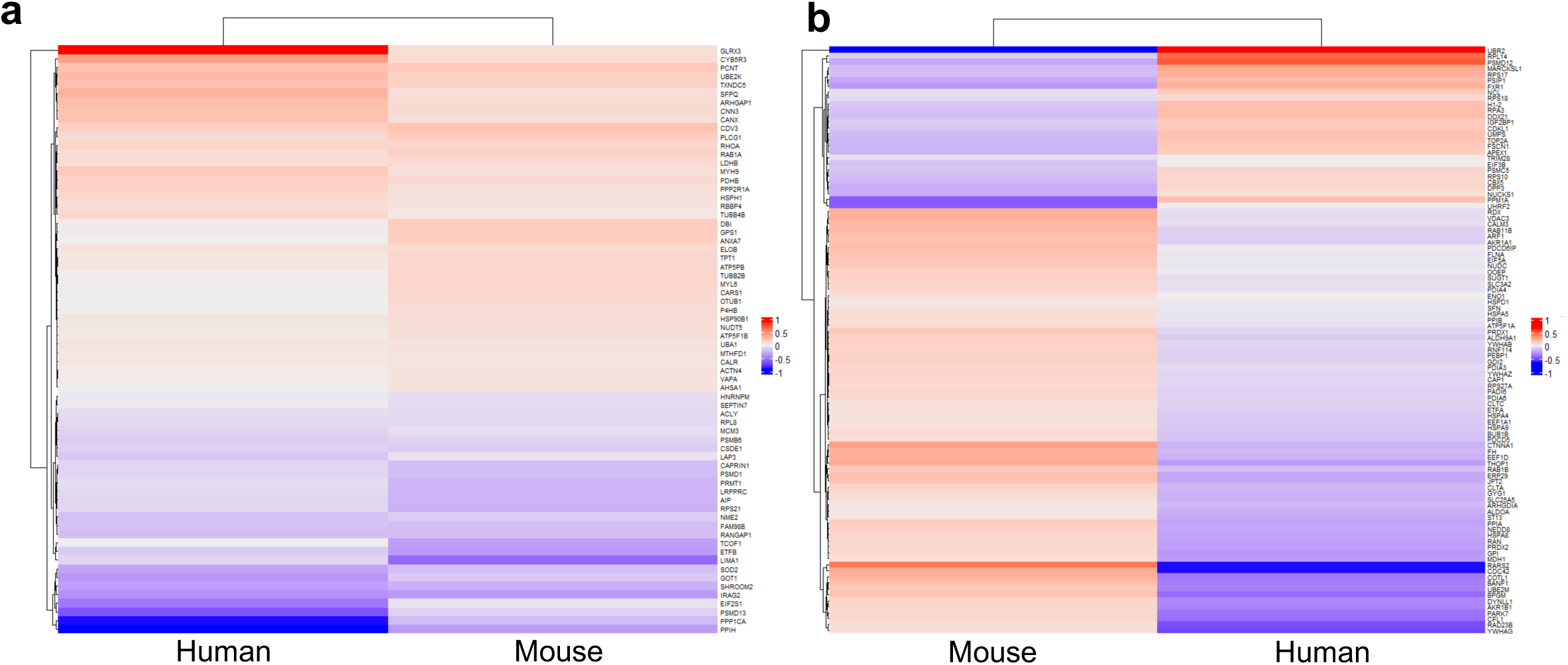
Further comparison of human 2-cell data to mouse and stem cell derivatives. **a, b** Heatmaps of median fold-change between alpha and beta blastomeres of proteins that were found to be significantly differential between alpha and beta clusters in the mouse data. The first heatmap corresponds to proteins that are changing in the same direction across the human and mouse blastomeres. The second heatmap illustrates proteins that have opposing median levels between human and mouse blastomeres.

## Methods

### Mouse embryo culture and sample collection

This research adhered to the regulations of the Animals (Scientific Procedures) Act 1986 - Amendment Regulations 2012 and was reviewed by the University of Cambridge Animal Welfare and Ethical Review Body. Experiments were approved by the UK Home Office.

Embryos were collected from 4-6 week old F1 females (C57BI6 x CBA, Charles River) following superovulation by injection of 5 IU of pregnant mares’ serum gonadotropin (PMSG, Intervet) and 5 IU of human chorionic gonadotropin (hCG, Intervet) 48 hrs later. Females were mated with F1 males (6 weeks - 52 weeks of age, C57BI6 x CBA, Charles River). Plugged females were culled by cervical dislocation to recover embryos at the required stage. Embryos, other than zygotes, were recovered in M2 medium (in house).

Zygote stage: zygote stage embryos (22 hrs post-hCG) were recovered in M2 medium with 1mg/ml of hyaluronidase (Sigma, H4272) in order to remove cumulus cells and subsequently washed through M2 medium without hyaluronidase. Samples were collected at 23-24 hrs post-hCG.

Early 2-cell stage: zygote stage embryos (29 hrs post-hCG) were recovered in hyaluronidase as above and subsequently cultured for 1-3h during division from the zygote to 2-cell stage. Following division to the 2-cell stage, samples were collected at 30-32 hrs post-hCG .

Late 2-cell stage: 2-cell stage embryos were recovered at 45 hrs post-hCG and samples collected at 46-48 hrs post-hCG.

4-cell stage: mid to late 2-cell stage embryos were recovered and one sister blastomere was microinjected as described below, to allow for identification of the 4-cell stage sisters originating from the injected 2-cell stage blastomere (mCherry positive pairs were distinguished by SCoPE2). The embryos were transferred to KSOM (Merck, MR-106-D), and live imaged during division from the 2- to 4-cell stage and collected at 55-57 hrs post-hCG. Division pattern, age post-division (2- to 4-cell stage) and division order were annotated for each embryo prior to collection. Uninjected and unimaged controls were also collected with the same timings.

The zona pellucida was removed prior to blastomere separation by brief acidic Tyrode’s solution treatment (Sigma, T1788), followed by washes in M2 media. Embryos were then transferred to 35mm petri dishes (Corning, 351008) coated in 1% agarose and covered with M2 media. Blastomeres were separated from each other using a thin glass capillary and transferred immediately to M2 medium. Separation took up to 1 min and had a survival rate greater than 80%. Zygotes were split in a similar manner to give rise to two intact split halves. Zygotes were split meridionally in alignment with the animal-vegetal axis as defined by the position of the polar body. After the embryos had been split, individual blastomeres or zygote halves were washed through 7-10 washes of PBS (Life Technologies, 10010056) followed by 5 washes in pure water (Optima LC/MS Grade, Fisher Scientific, W6500), before being finally resuspended in 1µl of water and transferred to individual wells of a 384 well plate (ThermoFisher, AB1384), on a cold block. In order to minimize sample contamination all surfaces were cleaned and filter tips used. Wash drops were not reused more than 8 times and were changed if a blastomere lysed. Two different glass pipettes were used for the PBS and water washes to prevent carry over. In each experiment, sample collection took up to 1 hr and plates were subsequently sealed with foil (ThermoFisher, AB0626).

For PCNA live imaging experiments, zygotes were recovered as described above and microinjected. Following microinjection, embryos were cultured in KSOM media (Merck, MR-106-D) under mineral oil (Biocare Europe) at 5% CO_2_, and 37°C overnight and allowed to cleave to the 2-cell stage. Prior to imaging, 2-cell embryos were split as above and transferred to an imaging dish. In experiments where further culture was required following imaging, split embryos were transferred to Global Total culture medium (LifeGlobal group, H5GT-030) under mineral oil (Biocare Europe) at 5% CO_2_, and 37°C, for 72 hrs.

For split embryo culture to the blastocyst stage, 2-cell stage embryos were split as above and single blastomeres cultured in drops of Global Total culture medium^88^ (LifeGlobal group, H5GT-030) under mineral oil (Biocare Europe) at 5% CO_2_, and 37°C, for 72 hrs.

For knockdown and overexpression experiments, zygote or 2-cell stage embryos were recovered as described above and microinjection performed. Following microinjection embryos were cultured in KSOM media under mineral oil at 5% CO_2_, and 37°C, until the required stage.

### dsRNA and mRNA synthesis

The mCherry sequence was amplified via PCR from the pRN3P-H2B-mCherry vector and cloned into the pRN3p vector via EcoRI/BamHI digestion (ThermoFisher, FD0054 and FD0274) and T4 ligation (New England Biolabs, M0202S). pRN3P-mCherry was linearised using KpnI (New England Biolabs, R3142S). In vitro transcription was carried out using the mMessage mMachine T3 kit (Thermo Fisher, AM1348) and purified via lithium chloride precipitation, according to the manufacturer’s instructions. pRN3P-Gap43-RFP and pRN3P-PCNA-Clover were linearised using SfiI (ThermoFisher, FD1824) and mRNA synthesized via in vitro transcription using T3 as above.

dsRNAs of 350-500bp length, were designed using the E-RNAi platform (Horn and Boutros, 2010) and amplified from mouse liver cDNA. In vitro transcription was carried out using the MEGAscript T7 kit (Thermo Fisher, AM1334) and purified via lithium chloride precipitation, according to the manufacturer’s instructions.

For overexpression experiments, Nedd8 and Gps1 sequences were amplified via PCR from mouse liver cDNA, and restriction sites and an HA tag added via PCR. Sequences were cloned in the pRN3p vector via EcoRI/BamHI (ThermoFisher, FD0274 and FD0054) and HindIII/BamHI (ThermoFisher, FD0505 and FD0054) digestion for Nedd8-HA and Gps1-HA respectively, followed by T4 ligation (New England Biolabs, M0202S). pRN3P-Nedd8-HA and pRN3P-Gps1-HA were linearised using SdaI (ThermoFisher, FD1194).

In vitro transcription using T3 was carried out as above. Primers used for dsRNA and mRNA synthesis are listed in Extended Data Table 5.

### Microinjection

Microinjection was performed as previously described^79^. Briefly, embryos were placed in a depression on a glass slide in M2 medium covered with mineral oil (Biocare Europe, 9305). Microinjection was performed using an Eppendorf Femtojet Microinjector with negative capacitance to facilitate membrane entry. Synthetic mCherry mRNA and Gap43-RFP were injected at a concentration of 200 ng/μl. For PCNA live imaging experiments, mRNA was injected at 100 ng/µl^83^ . For knockdown experiments dsRNAs were injected at a concentration of 1000 ng/μl. For overexpression, mRNA was injected at the indicated concentration (50 ng/μl or 500 ng/μl).

### qRT-PCR

RNA was collected from embryos at 48 hrs post-injection, using the Arcturus PicoPure RNA isolation kit (Arcturus Bioscience, KIT0204), according to the manufacturer’s instructions. Quantitative reverse transcriptase polymerase chain reaction (qRT-PCR) was performed using a StepOne Plus Real-time PCR machine (Applied Biosystem) and the Power SYBR Green RNA-to-CT 1-Step Kit (Life Technologies, 4389986). Relative mRNA expression levels of genes of interest were calculated using the ddCT method, with normalization to Gapdh. The primers used for qRT-PCR are listed in Extended Data Table 4.

### Immunofluorescence and confocal imaging

Blastocyst or 8-cell stage embryos were fixed in 4% PFA for 20 min at room temperature, and then washed through PBST (0.1% Tween 20 (Sigma Aldrich) in PBS) three times. Embryos were then permeabilized in 0.5% Triton X-100 (Sigma Aldrich) in PBS for 20 min at room temperature and washed through PBST again, before being transferred to blocking buffer (3% bovine serum albumin (SIgma Aldrich) in PBST) for 3 hrs at 4°C. Samples were then incubated with primary antibody mixes (diluted in blocking buffer) overnight at 4 °C. The next day, embryos were washed through PBST and incubated in secondary antibody mixes (1:500 in blocking buffer) with DAPI (Life Technologies, D3571, 1:1000 dilution, in PBST) for 2 hrs at room temperature. Finally, samples were washed through PBST following incubation with secondary antibodies and then imaged. Imaging was carried out on a SP5 or SP8 scanning confocal microscope (Leica) using the 63X or 40X oil objective.

Primary antibodies used: goat monoclonal anti Sox17 (R&D Systems, af1924, 1:200), mouse monoclonal anti Cdx2 (Launch Diagnostics, MU392-UC (Biogenex), 1:200), rat anti HA (Roche, 11867423001, 1:100), rabbit anti-Nanog (Abcam ab80892, 1:200) and rabbit anti RFP (Rockland, 600-401-379, 1:500)

Secondary antibodies used: Alexa Fluor 488 Donkey anti-Mouse, (ThermoFisher Scientific, A21202); Alexa Fluor 568 Donkey anti-Rabbit (ThermoFisher Scientific, A10042) and Alexa Fluor 647 Donkey anti-Goat (ThermoFisher Scientific, A21447). All secondaries were used at a 1:400 dilution.

### Image analysis and statistics

Images were processed with Fiji software^89^ (2012, https://imagej.net/software/fiji/) to assess cell number and lineage allocation. Cell numbers were counted manually, using the multi-point counter function. For HA tag staining, regions of interest were defined for the nucleus and cytoplasm at the midplane of each cell and intensity was measured using the ImageJ measure function. HA tag intensity was normalised to DAPI intensity.

The statistical test used is indicated in the corresponding figure legend. In all cases, the two-tailed version of the test was used. Normality of the data was assessed using the Shapiro Wilk test. Statistical analysis was performed using Prism software (version 8, GraphPad, https://www.graphpad.com/scientific-software/prism/).

### Live Imaging

4-cell stage classification: Live imaging was performed with a SP5 scanning confocal microscope (Leica) using the 63x oil objective. 2- to 4-cell mouse embryos were imaged on glass-bottom dishes (MatTek, P35G-1.5-14-C) within a nylon mesh (Plastok) in KSOM media under mineral oil and kept in a humidified chamber at 5% CO_2_, and 37°C throughout imaging. Images were captured every 15 mins with a z-step size of 5μm. Time lapse recordings were processed with Fiji software to assess division order, division timing and division pattern.

PCNA cell cycle assessment: Imaging was performed with a spinning disk confocal microscope (3i Intelligent Imaging Innovations) using the 63x oil objective, from 36 to 46 hrs post-hCG during the transition from S to G2 phase. Single blastomeres from split 2- cell embryos were imaged on glass bottom dishes, within a nylon mesh in KSOM media under mineral oil, at 5% CO_2_, and 37°C. The imaging interval was 15 mins and the z-step size 5μm. After imaging a total of 30 single 2-cell stage blastomeres (15 split embryos) were collected for subsequent MS analysis as above. In a separate set of experiments blastomeres were allowed to undergo the second cleavage division during imaging and then removed from the imaging chamber and cultured to the blastocyst stage. Images were exported from SlideBook (3i Intelligent Imaging Innovations) and subsequently processed with Fiji software to assess S phase exit and division order when possible.

### Mouse stem cell culture and sample collection

CAG-GFP/tetO-H2B-mCherry mouse embryonic stem cells (ESCs) were used as carrier samples for the SCoPE2. Cells were cultured on gelatin coated plates at 5% CO2, and 37°C in N2B27 2iLIF media. N2B27 2iLIF was comprised of 50% Neurobasal-A (Gibco, 10888022), 50% DMEM/F-12 (Gibco, 21331020), 0.5% N2 (in-house), 1% B27 (Gibco, 10889038), 2mM GlutaMAX (Gibco, 35050038), 0.1mM 2-mercaptoethanol (Gibco, 31350010) and 1% penicillin/streptomycin (Gibco, 15140122), with 3mM CHIR99021 (Cambridge Stem Cell Institute), 1mM PD0325901 (Cambridge Stem Cell Institute) and 10 ng ml-1 leukaemia inhibitory factor (Cambridge Stem Cell Institute) supplemented.

To induce H2B-mCherry expression, CAG-GFP/tetO-H2B-mCherry ESCs were treated with Doxycycline (1 mg/mL) (Sigma-Aldrich, D9891-5G) for 6 hrs prior to collection.

ESCs were routinely passaged at 70% confluency following trypsinisation (Trypsin-EDTA 0.05%, Life Technologies, 25300054) for 4 minutes at 37°C. Feeder cell media was added to terminate the trypsinization and cells were dissociated by gentle pipetting and centrifuged for 4 minutes at 1000 rpm, before being re-plated at a 1:10 or 1:20 dilution. Feeder cell medium contained Dulbecco’s modified essential medium (Gibco, 41966052), 15% fetal bovine serum (Cambridge Stem Cell Institute), 1mM sodium pyruvate, 2mM GlutaMAX, 1% MEM non-essential amino acids (Gibco, 11140035), 0.1mM 2- mercaptoethanol and 1% penicillin/streptomycin. Cells were routinely tested for mycoplasma contamination.

For sample collection, cells were trypsinized as above, and resuspended in PBS (Life Technologies, 10010056). The cells were then pelleted by centrifugation for 4 minutes at 1000 rpm before a second PBS wash. A haemocytometer was then used to estimate cell density and cells were pelleted as above, before being finally resuspended in pure water at a density of 2000 cells/ul. 200,000-300,000 cells total were collected in 0.2ml PCR tubes (Starlab, A1402-3700) and stored at -80°C.

### Human stem cell culture and sample collection

The use of human ESCs (hESCs) was approved by the UK Stem Cell Bank Steering Committee and experiments complied with the UK Code of Practice for the Use of Human Stem Cell Lines. RUES2 hESCs (kindly provided by Ali Brivanlou) were used as carrier samples for SCoPE2. All cells were routinely tested for mycoplasma contamination.

RUES2 hESCs were cultured in a humidified incubator at 37°C and 5% CO_2_ in mTeSR1 (StemCell Technologies, 85850) on growth factor-reduced Matrigel-coated (Corning, 353046). For Matrigel coating, plates were incubated with 0.16mg/ml Matrigel in DMEM/F12 (Gibco, 21331020) at 37°C for 1 hour. Media was changed daily.

hESCs were routinely passaged every 4–5 days by dissociating in Accutase (ThermoFisher Scientific, A1110501) for 3 minutes at 37°C. Cells were collected in DMEM/F12 and centrifuged for 3 minutes at 1000 rpm before being re-plated in mTesR1 medium supplemented with 10μM ROCK inhibitor Y-27632 (StemCell Technologies, 72304) for 24 hours.

For sample collection, cells were dissociated as above for routine passage. RUES2 hESCs were resuspended in PBS. Cells were pelleted by centrifugation for 4 minutes at 1200 rpm before a second PBS wash. Cells were resuspended in pure water at a density of 2000 cells/ul. 200,000-300,000 cells total were collected in 0.2ml PCR tubes and stored at -80°C.

### Ethics statement

Human embryo samples for this study were collected in two different institutes: the University of Cambridge (United Kingdom), and the California Institute of Technology (United States). All the work complies with The International Society for Stem Cell Research (ISSCR) guidelines^90^.

Human embryo work at the University of Cambridge was performed in accordance with Human Fertility and Embryology Authority (HFEA) regulations (license reference R0193).

Ethical approval for the work was obtained from the ‘Human Biology Research Ethics Committee’ at the University of Cambridge (reference HBREC.2017.21). Informed consent was obtained from all participants in the study which included patients from the CARE Fertility Group and Herts & Essex fertility clinics. Supernumerary embryos were donated upon completion of IVF treatment. Patients were informed about the specific objectives of the project, and the conditions that apply within the license, before giving consent. Patients were also offered counseling and did not receive any financial inducements for their donation.

Human embryo work at the California Institute of Technology was approved by the California Institute of Technology Committee for the Protection of Human Subjects (IRB number 19–0948). Human embryos at the two pronuclei stage were obtained from the University of Southern California (USC) through the pre-existing USC IRB-approved Biospecimen Repository for Reproductive Research (HS-15–00859) after appropriate approval was obtained unanimously from the Biorepository Ethics Committee. Supernumerary embryos were donated upon completion of IVF treated from USC Fertility. Patients were informed of the general conditions of the donation, as well as the objectives, and methodology of human embryo research. They were offered counseling and alternatives to donation, including discarding embryos and continued cryopreservation of embryos. Patients were informed that they would not benefit directly from the donation of embryos to research.

### Human embryo culture and sample collection

#### University of Cambridge

A total of 8 donated two pronuclei stage human zygotes (day 1 post fertilization) from two patients were used for this study. Embryos were warmed and cultured according to the above regulations. Cryopreserved day 1 embryos were thawed with the Origio thaw kit (REF10984010A) following the manufacturer’s instructions. Briefly, the Global Total human embryo culture medium (LifeGlobal group, H5GT-030) was incubated at 37°C and 5% CO_2_ overnight before thawing. The next day, the straw containing the embryo was immersed in prewarmed (37°C) water for 1 min. The embryo was then transferred into vial 1 (5min), vial 2 (5 min), vial 3 (10 min), and finally into vial 4. Thawed embryos were finally incubated in drops of the pre-equilibrated Global Total human embryo culture medium under mineral oil (Irvine Scientific, 9305). Embryos were incubated for a total of 12 hrs overnight at 37°C,, and 5% CO_2._ The following day the zona pellucida of the 2-cell stage human embryos was removed by brief acidic Tyrode’s solution treatment (Sigma, T1788).

Embryos were then bisected and single blastomeres transferred to M2 media before being washed and processed as above into 384-well PCR plates (ThermoFisher AB1384) in 1μl of pure water (Optima LC/MS Grade, Fisher Scientific W6500).

Of 8 two pronuclei zygote stage human embryos, 7 embryos developed to the 2-cell stage for sample collection and 5 were included in the final analysis.

#### California Institute of Technology

A total of 22 donated two pronuclei stage human zygotes (day 1 post fertilization) from 5 patients were used for this study. Cryopreserved day 1 embryos were warmed with the Embryo Thaw Media Kit following the manufacturer’s instructions (Fujifilm Irvine Scientific, 90124). Briefly, the Continuous Single Culture-NX Complete medium (Fujifilm Irvine Scientific, 90168) was incubated at 37°C and 5% CO_2_ overnight before thawing. The next day, the straw containing the embryo was defrosted at room temperature for 30 s and immersed in prewarmed (37 °C) water for 1 min until the ice melted. The embryo was then transferred into T-1 (5 min), T-2 (5 min) and T-3 (10 min) solutions for slow thawing, before being finally transferred to Multipurpose Handling Medium (Fujifilm Irvine Scientific, 90163) for recovery. Thawed embryos were then incubated in drops of pre-equilibrated Continuous Single Culture-NX Complete medium under mineral oil (Irvine Scientific, 9305). Embryos were incubated at 37°C, and 5% CO_2_ for 6-12 hrs until they reached the 2-cell stage.

Assisted hatching was performed using laser pulses at 200 μs (Lykos laser: Hamilton Thorne, Beverly, MA, USA). Embryo Biopsy Medium (Irvine Scientific, 90103) was then used to separate the blastomeres from each other, followed by gentle pipetting of the embryo to remove it from the zona pellucida and isolate individual sister blastomeres. Blastomeres were then washed and processed as above into 384-well PCR plates (ThermoFisher, AB1384) in 1μl of pure water (Optima LC/MS Grade, Fisher Scientific, W6500).

Of 22 two pronuclei zygote human embryos, 20 embryos developed to the 2-cell stage for sample collection and 8 were included in the final analysis.

### Single-cell RNA sequencing analysis

Single-cell RNA-sequencing analysis was performed on early, mid, and late 2-cell stage blastomeres using the previously published dataset by Deng *et al.*^91^ (GSE45719). Reads were aligned against the reference genome GRCm39 and count matrices were made using kallisto | bustools^92^. Further downstream analyses were performed in Python using the Scanpy toolkit (version 1.9.1) and Anndata (version 0.8.0)^93^. None of the cells were filtered for mitochondrial or ribosomal content. Initial analysis including normalization, scaling, and identification of highly variable genes was performed using the Scanpy preprocessing toolkit. Single-cell data was further visualized using several Scanpy plotting tools including: sc.pl.umap, sc.pl.clustermap, and sc.pl.heatmap.

### Mass spectrometry (MS) sample preparation

#### SCoPE2 and pSCoPE sample preparation

##### Isobaric carrier & reference

ample preparation and analysis was performed as described by Petelski et al.^46^. Briefly, mouse embryonic stem cells at a density of 2,000 cells/μl in 100μl water were lysed through the mPOP method (frozen cells were subjected to a rapid heat cycle of 90°C for 10 minutes, and then were cooled to 12°C)^13,46^. Trypsin Gold (TG, Thermo) and triethylammonium bicarbonate (TEAB, pH = 8, Sigma) were added to the cell lysate to final concentrations of 10 ng/μl and 100 mM, respectively. The sample, once mixed with TG and TEAB, was subjected to 37°C overnight (16-18 hours) to digest proteins into peptides. To ensure adequate miscleavage rate (<20%), a small amount (1μl) of the cell digest was evaluated via LC-MS/MS. The cell digest was split into two samples, one for the carrier and the other for the reference. The carrier was labeled with TMT 126 and the reference was labeled with TMT 127N, with the labeling reaction proceeding for 1 hour. The reaction was quenched with 1% hydroxylamine (HA) for 30 minutes. Labeled material was then evaluated via LC-MS/MS for labeling efficiency (> 99%). Carrier and reference materials were kept frozen -80°C until needed for multiplexing with single cells.

##### Single blastomere cells and half zygotes

Frozen blastomeres that were collected in a 384-well plate were lysed by rapidly heating in a thermocycler to 90°C for 10 minutes and then cooled to 12°C. To each well (with a single blastomere or a water serving as the control), TG and TEAB were added to the cell lysate to final concentrations of 10 ng/μl and 100 mM, respectively. The plate was then subjected to 37°C for three hours. Each well then received 0.5μl of selected TMT reagents and the plate was incubated at room temperature for 1 hour. The labeling reaction was quenched with 0.5μl of 1% HA at room temperature for 30 minutes. Single blastomeres were then combined with 200 carrier and 5 reference cells to form a TMT set (Extended Data Fig. 1c). Each TMT set was dried down and resuspended in 1.1 μl of HPLC-grade water.

### Sample preparation for label-free mass-spec single-cell proteomics

A total of eight 2-cell stage human embryos were used for label-free mass-spectrometry analysis. Frozen blastomeres that were collected in a 384-well plate were lysed by rapidly heating to 90°C for 10 minutes and then cooling to 12°C. To each well (with a single blastomere or a water serving as the control), TG and TEAB were added to the cell lysate to final concentrations of 10 ng/μl and 100 mM, respectively. The plate was then kept at 37°C for 3 hours to facilitate protein digestion. Each blastomere was dried down in a speed-vac and resuspended in 1.1 μl of HPLC-grade water for subsequent mass-spectrometry acquisition.

### Mass spectrometry acquisition methods

TMT sets of single blastomeres were analyzed according to the SCoPE2 protocol guidelines. Specifically, 1 μl out of 1.2 μl of each SCoPE2 pooled sample was loaded onto a 25 cm × 75 1 μm IonOpticks Aurora Series UHPLC column (AUR2-25075C18A). Buffer A was 0.1% formic acid in water and buffer B was 0.1% formic acid in 80% acetonitrile / 20% water. A constant flow rate of 200 nl/min was used throughout sample loading and separation. Samples were loaded onto the column for 20 min at 1% B buffer, then ramped to 5% B buffer over 2 min. The active gradient then ramped from 5% B buffer to 25% B buffer over 53 min. The gradient then ramped to 95% B buffer over 2 min and stayed at that level for 3 min. The gradient then dropped to 1% B buffer over 0.1 min and stayed at that level for 4.9 min. The total run time of each sample took 95 minutes total. All samples were analyzed by a Thermo Scientific Q-Exactive mass spectrometer from minutes 20 to 95 of the LC loading and separation process. Electrospray voltage was set to 2200 V and applied at the end of the analytical column. To reduce atmospheric background ions and enhance the peptide signal-to-noise ratio, an Active Background Ion Reduction Device (ABIRD, by ESI Source Solutions, LLC, Woburn MA, USA) was used at the nanospray interface. The temperature of the ion transfer tube was 250 °C, and the S-lens RF level was set to 80.

### Analysis of raw SCoPE2 and pSCoPE MS Data

Raw data were searched by MaxQuant (version 1.6.17) against a protein sequence database including all entries from the mouse or human SwissProt database (depending on which samples were being analyzed) and known contaminants such as human keratins and common lab contaminants (default MaxQuant contaminant list).

Within the MaxQuant search, we specified trypsin digestion and allowed for up to two missed cleavages for peptides having from 7 to 25 amino acids. Tandem mass tags (TMTPro 16plex) were specified as fixed modifications. Methionine oxidation (+ 15.99492 Da), and protein N-terminal acetylation (+ 42.01056 Da) were set as variable modifications. Second peptide identification was disabled. Calculate peak properties was enabled. All peptide spectrum matches (PSMs) and peptides found by MaxQuant were exported in the evidence.txt files. These evidence files were then analyzed together by DART-ID^94^. The data from the files processed from DART-ID^94^ was then processed with the SCoPE2 pipeline^13^ with minor modifications, with filtering parameters including PEP < 0.03 and PIF > 0.8. Reverse matches and contaminants were also removed.

SCoPE2 pipeline is available here: (https://zenodo.org/record/4339954#.YnHcYSfMLOQ)

The carrier and reference material in these experiments are clearly a different cell type from the single cells that we have processed. Obtaining a large enough number of blastomeres to use as carrier and reference material for all the SCoPE2 sets was not possible. Although the cell types are different, we were still able to sequence and quantify many peptides that were heavily enriched in the blastomeres as shown in Fig. 2c. These peptides and the corresponding proteins include classical markers of blastomeres and are biologically relevant, as evidenced by our analyses of mouse ESCs carriers vs blastomeres. The protein differences associated with alpha-beta asymmetry are also observed in our label free DIA experiments that did not use a carrier, suggesting that the choice of carrier is unlikely to strongly influence our results.

### Label-free DIA analysis and DIA-NN search parameters

Some human blastomeres were analyzed by label-free DIA using a 100 minute total gradient, of which 63 minutes were active (12 to 75 minutes). More specifically, the gradient used is as follows: 4% buffer B (minutes 0 - 11.5), 4%-8% buffer B (minutes 11.5 - 12), 8%-35% buffer B (minutes 12 - 75), 35%-95% buffer B (minutes 75-77), 95% buffer B (minutes 77 - 80), 95%-4% buffer B (minutes 80 - 80.1), 4% buffer B (minutes 80.1 - 100). Each duty cycle consisted of 2 MS1 windows with ranges from 480 - 1500 m/z.. Each MS1 was followed by 3 MS2 windows spanning its m/z range (2x 1 MS1 full scan x 3 MS2 windows). The size of the 6 MS2 windows in each duty cycle were variable and were as follows: 480 - 530 m/z; 530 - 590 m/z; 590 - 650 m/z; 650 - 750 m/z; 750 - 1000 m/z; 1000 - 1500 m/z. Each MS1 and MS2 scan was conducted at 140k resolving power, 3×10^6^ AGC maximum, and 600 ms maximum injection time for both MS1 and MS2 scans.

Raw data was searched with DIA-NN (version 1.8)^95^ against a protein sequence database that included entries from the human SwissProt database (SwissProt_human_09042017, containing 20,218 proteins). The fragment sizes were set from 200 - 1800 m/z, with N-terminal methionine excision enabled. We specified the search for trypsin digestion and set the maximum number of missed cleavages to 1. Scan window radius was set 1, while the peptide lengths were set at 7 - 30 amino acids.

### K-means Clustering

To estimate the stability of the cell classification that was accomplished via k-means clustering, we computed the stability of cluster assignment. Through 200 iterations in which the starting cell centroid was changed for each cluster, we estimated the probability of cluster assignment for each. The overwhelming majority of cells have a high probability of landing in the same cluster consistently when initial conditions are changed. There are some blastomeres (n=9) that seem to exhibit unstable cluster assignment, which we have been unable to link to division order, division timing, or division pattern. For simplicity’s sake, we arbitrarily termed these clusters as alpha and beta. The same approach was used for both the human blastomeres from 2-cell embryos and the cut zygotes data.

### Determining differential proteins between alpha- and beta-cell types

Once cells were assigned to their respective classes via k-means clustering (k=2), we determined which proteins were significantly differentially abundant between the two groups of blastomeres using a series of Kruskal-Wallis tests (effectively a Mann-Whitney- Wilcoxon test). At least three observations per group were required for each protein. P values of the tested proteins were adjusted for multiple hypotheses through the BH method to estimate the false discovery rate (FDR). A threshold of 5% FDR was implemented as the cutoff for significance for all results.

From these analyses, we obtained a list of differentially abundant proteins in 2-cell embryos and used these proteins to plot two heatmaps in order to designate between the early and late 2-cell stages. The heat maps represent the proteins x blastomeres matrices. The columns of each heatmap were ordered by descending degree of asymmetry of sister blastomeres. The leftmost and rightmost columns correspond to blastomeres from the same embryo, a pattern that continues to the center of the heatmap.

Overall, 349 proteins that are distinguishing the alpha-beta clusters. Out of this list, 163 proteins were quantified in the mouse zygote data, which is 47% of the defined alpha- beta proteins.

Overall, we quantified an average of 3586 peptides mapping to 1043 proteins in the mouse blastomere samples, and a mean of 2895 peptides mapping to 759 proteins in the human blastomere samples.

### Comparison between bulk stem cells and mouse blastomeres

#### Blastomere Peptide Enrichment

We plotted the reporter ion intensities (without any data processing) of a representative blastomere and its respective carrier on the log10 scale. In doing so, we find that some peptides are much more abundant in one blastomere as compared to a 200-cell sample. In order to find what biological processes are enriched generally across mouse blastomeres as compared to ESCs, we obtained the precursor ratios of each blastomere to each respective ESC carrier. Then, we took the median across all blastomere-ESC pairs to obtain the median ratio for each precursor. Then, these ratios were further collapsed to the protein level by taking the median across all peptides mapping to that protein. With this list, we were able to rank proteins from greatest to least ratios, then input this ranked list into GOrilla using “single ranked list of genes” mode. From this output, we find that protein transport and protein degradation are largely enriched in blastomeres.

### Protein Set Enrichment Analysis (PSEA) for Alpha vs Beta Comparison

To determine which processes are differential between alpha- and beta- cell clusters, we first downloaded protein sets from MGI (MGI Data and Statistical Reports (jax.org)). These terms were filtered for proteins by Gene Symbols that were quantified in the mouse data. For each GO term, proteins by Gene Symbol that were associated with that GO term were collected into a single dataframe. That dataframe was further stratified into two groups: alpha- and beta- type. Each group was required to have greater than three observations. The two distributions per GO term were tested using the Kruskal Wallis test (effectively a Mann-Whitney-Wilcoxon test). P values of the tested GO terms were adjusted for multiple hypotheses, using the BH procedure to estimate the False Discovery Rate (FDR). GO terms were deemed significant if they passed the 5% FDR threshold.

For the mouse data, there were many GO terms that were significant (n = 2898 at 5% FDR, n = 1499 at 5% FDR and with greater than 2 proteins associated with the term). In order to make sense of all the terms, the data was stratified into themes of protein degradation, protein transport, translation, and metabolism through filtering of the names of the GO terms. These themes were further grouped into sub-themes in the same manner. This approach was also used for the human 2-cell stage data, using protein sets defined for human data.

### Ribosomal Protein Analysis

For each ribosomal protein (RP) that was quantified, we used the Mann-Whitney- Wilcoxon test to understand whether the abundance of the particular RP was different between alpha and beta cells across all available mouse blastomeres (from early 2-cell to 4-cell stage). From these analyses, we found that eight RPs are significantly differential (q-value < 0.05). The distribution of these proteins’ fold-changes between sister alpha and beta cells were plotted as boxplots at each stage.

### Vegetal cell analysis

To identify whether alpha / beta polarization is associated with vegetal cell identity, we clustered 4-cell stage blastomeres with well differentiated alpha-beta character based on their relative protein levels, as shown in Fig. 5e. As expected, the blastomeres clustered by alpha / beta polarization, and this clustering also portioned the vegetal cells. We evaluated the statistical significance of this portioning using the hypergeometric distribution to compute the cumulative probability (p-value) that the vegetal cells exhibit the observed association with alpha character or larger if sampled randomly.

### Across the stages Analysis

To assess which biological processes are driving this trend, fold changes between sister blastomeres assigned to opposing classes were calculated for each protein. With these values, we sought to identify functionally related proteins that covary among the three stages using spearman correlation analysis. For this analysis, we looped through each protein or each protein set and correlated the stages (set to be numerical) to fold changes between alpha and beta blastomeres of normalized protein abundances. From each correlation, we also obtained a p-value. Protein sets were required to have more than two proteins and more than 50% of proteins quantified. These p-values were then corrected for multiple hypotheses.

### Comparison between human and mouse 2-cell embryos

#### Pairwise Cell-to-Cell Correlation Heatmap

All proteins that were quantified in both datasets were used to calculate pairwise spearman correlations between human and mouse blastomeres at the 2-cell stage. The heatmap of spearman correlations was then plotted to have mouse blastomeres on the x-axis and human blastomeres on the y-axis. The human blastomeres are ordered in the same way as the dendrogram presented. The mouse blastomeres are clustered by their respective cluster-type, alpha or beta. We observe two distinct clusters, meaning that human 2-cell stage embryos also exhibit a similar proteome asymmetry.

#### Intersected Protein Set Heatmap

Protein sets that were found to be differentially abundant between alpha and beta cells in mouse were intersected with protein sets that were found to be differentially abundant between the respective two clusters. Both heatmaps were hierarchically clustered on the rows (protein sets) and blastomeres on the columns were clustered according to the cell classification via k-means clustering. Each tile in the heatmap represents the median value on the log2 scale of that particular protein set in a particular blastomere.

### Cut zygotes analysis

Upon normalization and performing k-means clustering in the same manner as in the mouse and human data, all quantified proteins in the zygotes were used to perform principal component analysis. Each zygote half fell into a cluster opposite its partner half, which we simply termed in this case “Cluster 1” and “Cluster 2”. The difference in the first principal component (PC1) values were taken for each zygote pair and plotted in a descending order as a barplot. Then, protein fold changes between the partner cut pieces were calculated for each zygote (these values were calculated consistently by finding the difference between the zygote piece in Cluster 1 and in Cluster 2. These vectors of protein fold changes for each zygote were then correlated pairwise to protein fold changes of each mouse 2-cell stage blastomere (which were consistently calculated as the difference between alpha and beta cells), resulting in a correlation matrix. These results were plotted as distributions per zygote, with the median of each distribution highlighted as a purple diamond (Fig. 3f).

In order to see the overall correlation between the zygote and mouse 2-cell blastomeres, the median fold change of each protein was calculated across all samples in respective groups (zygotes and 2-cell embryos). With these two vectors, a scatterplot of mouse 2- cell embryos protein fold-changes was plotted against the zygote fold-changes. The correlation of these vectors was positive (with a value of 0.44) and was highly significant (p-value = 1.39 x 10^-9^).

### Split Blastomere Experiment Analysis

Each blastomere that was used for MS analysis was prepared in a similar manner as was described in section titled “***Sample preparation for label-free mass-spec single-cell proteomics*** ” The MS acquisition was altered, so that peptides mapping to alpha-beta proteins were prioritized using prioritized SCoPE (pSCoPE)^81^, which was set up as described below, broadly following figure S4 from Huffman et al. 2023.

Gas phase fractionation (intensity-based quintiles spanning: 450-550, 550-623, 623-694, 694-788,788-1436 m/z respectively) was carried out to generate an empirical library using 5x TMT labeled mouse ESC carrier-reference runs. Post-acquisition, the runs were searched alongside all previous single cell DDA runs using Spectronaut (version 16.1), the generated spectral library was filtered at 5% FDR.

Subsequently, a 1x TMT labeled carrier reference sample was analyzed in DIA mode (using method outlined in Supplementary Table 5 from 10) to record accurate retention times for precursors. The run was searched using Spectronaut (filtered at 1% FDR).

The inclusion list was generated using peptides confidently identified in the 1x run. Peptides were split into 3 tiers, the highest tier contained peptides belonging to alpha and beta proteins, while the following two tiers contained peptides split by intensity, confidence of identification and precursor ion fraction. The inclusion list is provided as a supplementary file.

Prioritization was implemented using MaxQuant.Live Version 2.2.011. Targeting parameters were the same as in Supplementary table S13 (Method 5) of Huffman et al. 2023^80^, with the exception that survey scan life cycle was set to 1500ms, MS2 resolution was 70,000 and MS2 max injection time was 256ms.

The resulting data was searched using MaxQuant and then normalized as described previously in the section titled “***Analysis of raw SCoPE2 and pSCoPE MS Data*** ”, except that the final protein x samples matrix was normalized relative to the mean of all analyzed cells. Then, for each blastomere, the median abundance of alpha proteins was divided by the median abundance of beta proteins to estimate the alpha-beta fold change of the analyzed cell. By designating which proteins are alpha and which are beta from previous clustering the “likeness” or polarization of blastomeres could be calculated, with a higher median alpha protein level indicating alpha identity, and higher median beta protein level indicating beta identity. After calculating the alpha-beta protein fold change or alpha-beta likeness of a blastomere, it can be inferred that the cultured sister blastomere is of the opposite identity, as 2-cell stage sisters consistently separated into opposing clusters. Each blastomere’s fold change was then plotted against the proportion of epiblast cells in resultant blastocyst from the sister cell that was cultured. An overall positive trend is observed in the data, Fig. 4c. The distributions of alpha-beta fold changes were further analyzed by separating all blastomeres into two groups: (1) those whose sister blastomeres gave rise to blastocysts containing equal to or more than 4 epiblast cells and (2) those whose sister blastomeres gave rise to blastocysts with less than 4 epiblast cells, indicating the health of the embryo at this developmental stage.

---

**Extended Data Table 5:**
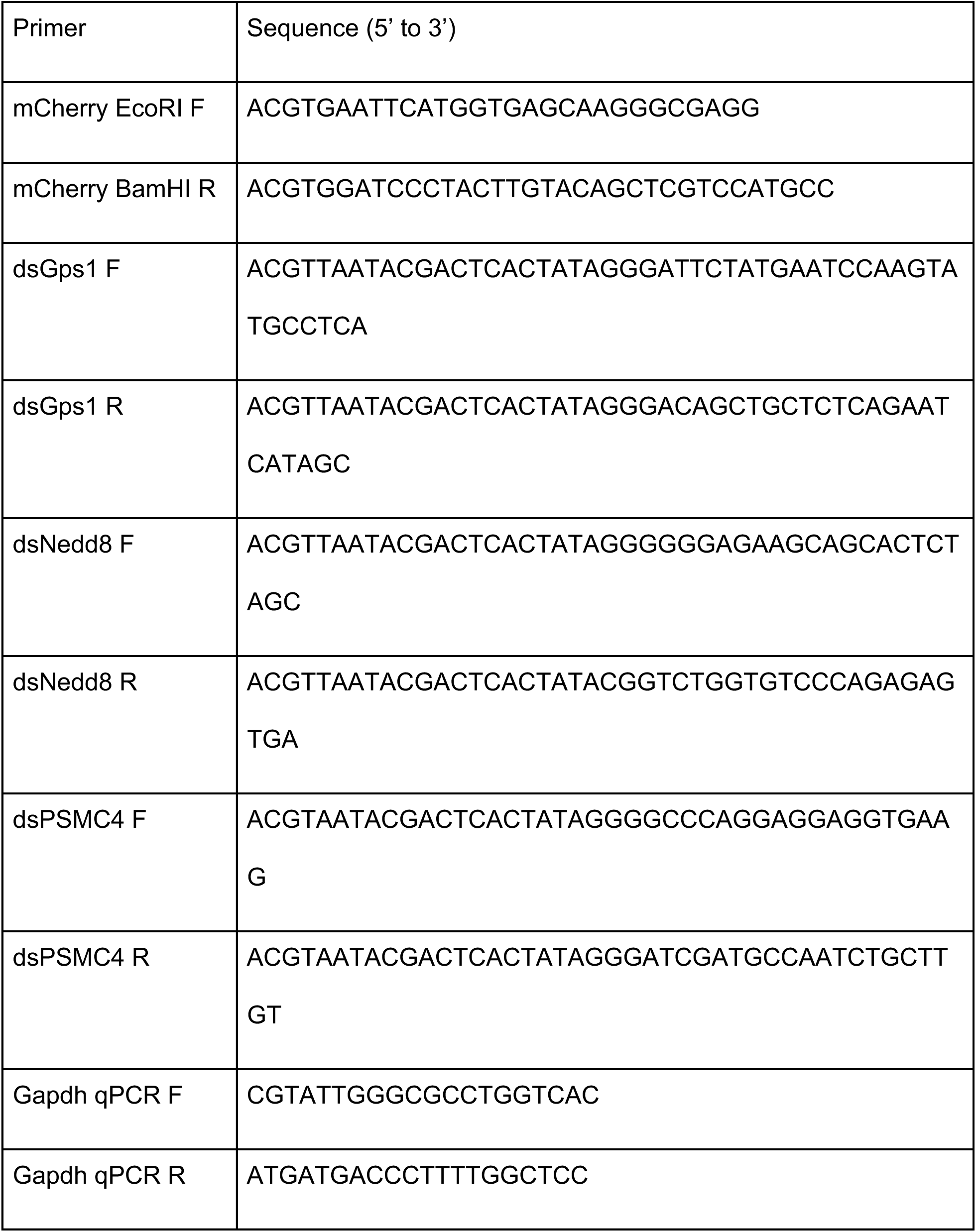
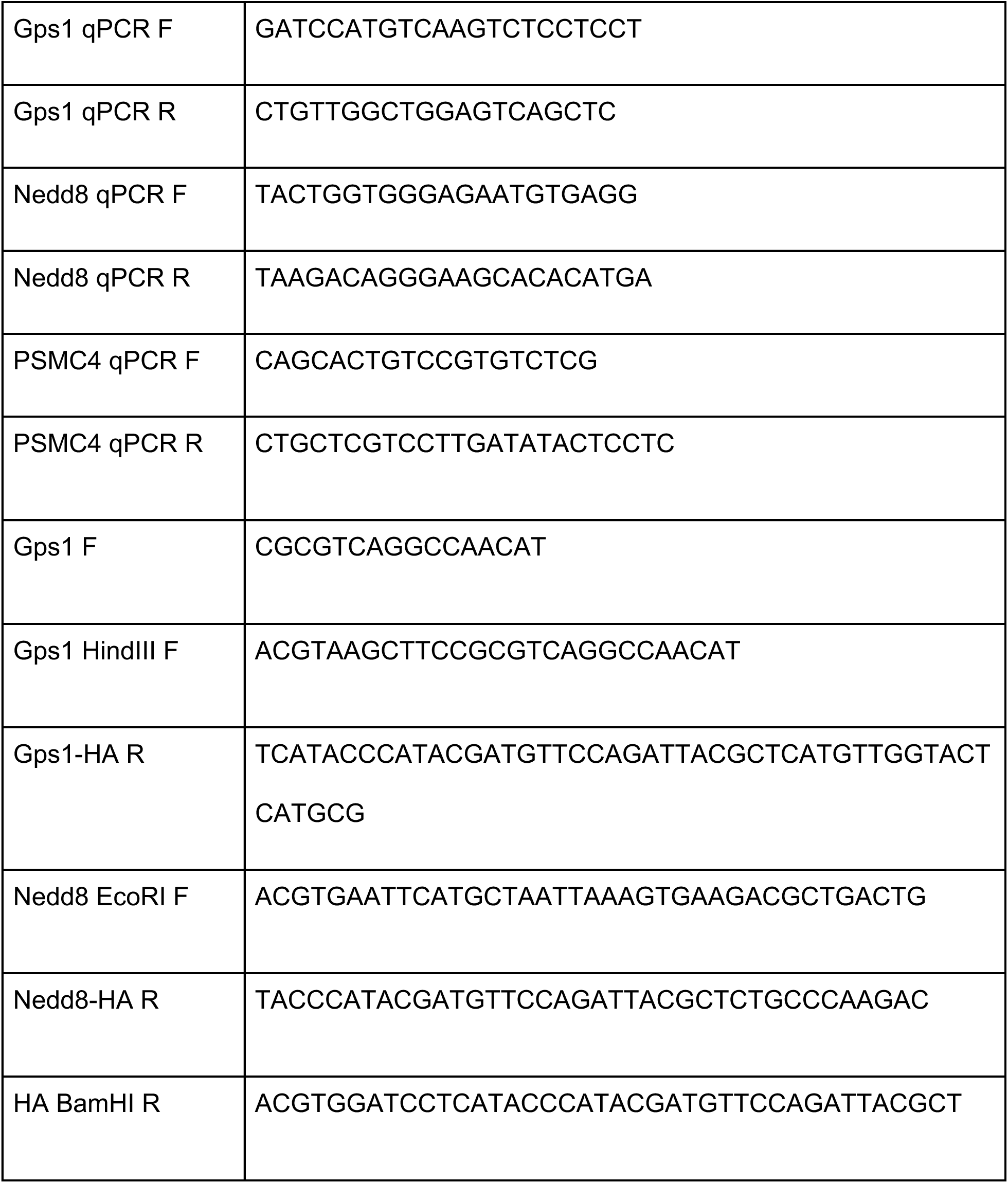
Primer sequences.

## Acknowledgements

We are thankful to the CARE Fertility Group, Herts & Essex fertility clinics and USC Fertility for their support. L.K.I-S is funded by the Rosetrees Trust (M877). M.M is supported by the Leverhulme Trust (RPG-2018-085). This work was funded by Wellcome Trust (207415/Z/17/Z), Open Philanthropy Grant, Weston Havens Foundations, and the NIH R01 (HD100456-01A1) grant (no NIH funding was used to support human embryo work) to M.Z-G., and by a New Innovator Award from the NIGMS from the National Institutes of Health to N.S. under Award Number DP2GM123497, an R01 award from NIGMS from the National Institutes of Health to N.S. under award number R01GM144967, and an Allen Distinguished Investigator award through the Paul G. Allen Frontiers Group to N.S.

## Author contributions

L.K.I-S and M.M performed embryo collection, experimental work on embryos, mouse and human stem cells, and data analysis. A.A.P. performed the mass spectrometry sample preparation, mass spectrometry maintenance, and data analysis. A.A.P. and N.S designed data analysis approaches. A.F. helped with functional experiments. H.S. and G.H. helped with data analysis and mass spectrometry maintenance. J.D. helped with the label-free DIA mass spectrometry acquisition and data analysis. A.A.P and S.K. performed pSCoPE analysis. V.J. performed single-cell RNA sequencing analysis. A.W., B.A.T. and C.W.G. performed the human embryo work at Cambridge. R.S.M., R.J.P., L.L., A.A., and E.S.V. performed human embryo work in California. A.A.P, L.K.I-S, N.S., and M.Z-G. wrote the manuscript. L.K.I-S, and M.Z-G conceived the project. N.S and M.Z-G. supervised this work.

## Data Availability

Metadata, raw data and processed data are organized according to community recommendations^96^ and are freely available at MassIVE: MSV000089353.

Direct Download Link: ftp://MSV000089353@massive.ucsd.edu Peer Reviewer Login: https://massive.ucsd.edu/ProteoSAFe/dataset.jsp?task=80e33c86f3034328ac5499c098dec6d9

## Code availability

All code was written in R (some of which was adapted from the SCoPE2 pipeline at https://github.com/SlavovLab/SCoPE2). The code is available as a collection of supplementary files in the MassIVE repository above.

